# Easy-to-use whole-genome sequencing workflows and standardized practices to uncover hidden genetic variation in *Synechocystis* PCC 6803 wild-type and knock-out strains

**DOI:** 10.64898/2026.04.08.717167

**Authors:** Marius Theune, Ruben Fritsche, Niklas Küppers, Marko Böhm, Petra Kolkhof, Florian Paul, Ovidiu Popa, Ellen Oldenburg, Anika Wiegard, Ilka M. Axmann, Kirstin Gutekunst

**Affiliations:** Molecular Plant Physiology, Bioenergetics in Photoautotrophs, Institute for Biology, University of Kassel, D-34132 Kassel, Germany; Institute for Synthetic Microbiology, Biology Department, Heinrich Heine University Düsseldorf, D-40225, Düsseldorf, Germany; Alfred Wegener Institute Helmholtz Center for Polar and Marine Research, Bremerhaven, Germany; Institute of Quantitative and Theoretical Biology, Biology Department, Heinrich Heine University Düsseldorf, D-40225, Düsseldorf, Germany; Cluster of Excellence on Plant Science, Heinrich Heine University Düsseldorf, Düsseldorf, Germany

## Abstract

Knock-out mutants are often used to study gene function by disrupting a specific gene and comparing the mutant to a wild-type strain. Reliable interpretation, however, requires that the two strains differ only by the intended mutation and that the observed phenotype is caused specifically by the deleted gene. In the highly polyploid cyanobacterium *Synechocystis* sp. PCC 6803, this is particularly challenging because incomplete segregation can mask genetic heterogeneity or secondary suppressor mutations. The genetic variation among laboratory wild-type lines can further confound phenotypic analyses. We show that these challenges can be addressed by routine strain validation via whole-genome sequencing (WGS). To this end, we developed and tested user friendly workflows for short-read (Illumina), long-read (Oxford Nanopore Technologies; ONT), and hybrid data, providing standardized quality control, variant calling, and structural variant detection. We benchmarked their performance in detecting single-nucleotide polymorphisms (SNPs), small indels, and structural variants using simulated datasets across different coverages and mixed populations. Applying the workflows to three *Synechocystis* sp. PCC 6803 wild-type lines revealed multiple sequence and structural differences relative to the reference genome, including previously undescribed genetic variants, underscoring the importance of documenting the strain background and the value of long-read sequencing. Characterization of two independent 6-phosphogluconate dehydrogenase (*gnd*) knock-out mutants and their complemented strains highlighted how a failed rescue can reveal a phenotype unrelated to the intended knock-out. An automated literature analysis revealed that only a minority of the investigated *Synechocystis* studies that used knock-out mutants included complementation as a control (39%), whereas this practice is more common in studies involving *Escherichia coli* (63%) and *Saccharomyces cerevisiae* (55%). Based on these results, we propose a practical guide for standardizing knock-out phenotyping in *Synechocystis* PCC 6803. Combined with accessible workflows for routine whole-genome validation, this framework aims to support more robust and reproducible knock-out studies in the future.

## Introduction

The unicellular cyanobacterium *Synechocystis* sp. PCC 6803 (hereafter *Synechocystis*) was one of the first organisms to be fully sequenced (1) and is widely used as a model organism. Its efficient integration of foreign DNA into the genome via homologous recombination and suitability for genetic engineering have made it a preferred model organism for basic research and green biotechnology applications (2–4). To investigate a gene’s function, knock-out mutants are often employed, in which the gene of interest is disrupted or replaced, and the resulting strain is compared with a wild-type strain. A deviant phenotype is then used to elucidate a gene’s function. While creating knock-out mutants is straightforward, *Synechocystis* presents a significant challenge for such genetic studies: its polyploid genome. (5). Variable copy numbers across cells and growth conditions necessitate segregation periods after genetic manipulation (6,7). While polyploidy can be beneficial when studying non-segregated knock-out strains of essential genes, the slow segregation increases the risk of secondary mutations, particularly suppressor mutations, arising under selection pressure from the intended genomic change (8,9). Controlling for secondary mutations or identifying strain-specific ones is particularly important, given the well-documented genetic diversity even among *Synechocystis* wild-type strains (10–12). These variations are usually detected using next-generation sequencing techniques, such as Illumina sequencing, and by mapping the generated reads to a reference sequence. In this technique, genomic DNA is fragmented and bridge-amplified to create paired local-amplified sequences on a chip, which are then sequenced in parallel using a cyclic reversible-termination approach. It thereby sequences entire bacterial genomes with several hundred-fold coverage at a very low per-base-pair error rate, but is limited by the short read length of around 150 bp (13). Having only short reads poses major challenges for detecting larger structural variants or sequencing repetitive or non-unique genomic regions. Several techniques have been developed to overcome this limitation; a prominent one is Oxford Nanopore (ONT) sequencing (14). This technique fundamentally differs from polymerase-based approaches in that it sequences a long DNA strand by translocating it through a pore while measuring sequence-specific current shifts. This technique generates ultra-long sequencing reads, yielding high coverage; however, the per-base-pair error rate remains significantly higher (15). Since both techniques are commercially available, bacterial WGS has become accessible and affordable, even for non-specialized laboratories (16–18). However, filtering and analyzing the generated data is labor-intensive, potentially preventing laboratories from routinely checking their mutant or wild-type strains (19). An alternative approach to determining whether a phenotype is directly linked to a gene knock-out rather than to a secondary mutation is to generate a complemented strain by reintroducing the previously knocked-out gene. These controls are performed by either introducing the gene back to its original locus (complementation in a narrow sense) or to another genomic location or to an extrachromosomal plasmid, after which it is investigated whether the knock-out phenotype is rescued (20). A suitable strain for comparing different complementation strategies is the previously published 6-phosphogluconate dehydrogenase (gnd) knock-out mutant (21). As Gnd is a key enzyme of the oxidative pentose phosphate pathway (OPP), it is essential for light-activated heterotrophic growth (hereafter, heterotrophic growth) (22). It was recently reported that this strain contains a secondary frameshift mutation at position 399 of the glucose-6-P. dehydrogenase (zwf) gene, which was detected by targeted Sanger sequencing (23). This frameshift introduces an early stop codon, thereby truncating the protein and abolishing Zwf enzyme activity. Consequently, this strain was renamed Δgnd_zwf^399+A^. Zwf is another key enzyme of the OPP pathway, upstream of Gnd, and its knock-out mutant is also unable to grow under heterotrophic conditions (24). We created an independent *gnd* knock-out strain *(Δgnd_3)* in the same wild-type and complemented it using different techniques. We compare the different complementation strategies and show how a failed rescue in a complemented strain can reveal a phenotype unrelated to the intended knock-out. To facilitate the use of WGS, we created and validated publicly available, easy-to-use pipelines for genomic variant calling from paired Illumina and ultra-long-read ONT sequencing data, and used them to discover novel variants across different wild-type strains. Finally, we developed a guide that outlines many key steps to make *Synechocystis* knock-out studies as robust and reproducible as possible.

## Results

### Design of user-friendly, Galaxy-based pipelines for whole-genome sequencing (WGS) analysis of *Synechocystis* strains

To make WGS more accessible to non-specialized users, we created data analysis pipelines on a publicly available bioinformatics server (usegalaxy.eu, The Galaxy Community, 2024). Galaxy is free to use, provides sufficient computational power, includes a large set of preinstalled tools, and offers a graphical user interface that enables easy data management and processing. Using only tools available on the server, we created two pipelines that process either paired-end short next-generation sequencing reads (e.g., Illumina), long ONT sequencing reads, or a combination of both to detect mutations by mapping reads to the provided reference genome (S3 File). The pipelines handle all necessary steps, such as quality control, filtering, adapter trimming, mapping, and, finally, calling mutations and annotating them (Fig 1; more detailed overview in S1 Fig; detailed manual in S4 Text). Important settings for filtering sequencing reads and distinguishing true variants from false-positive calls are available in the Galaxy server’s graphical user interface, where they can be easily adjusted, with recommended default values provided. To increase sensitivity, two different callers were used in parallel when possible, and their results were combined. The pipelines return the called mutations and structural variations separately, in both a “.vcf” (variant call format) file for further analysis and a “.csv” (comma-separated values) file. These .csv files can be imported into Microsoft Excel using the provided macro (S5 Text), which creates a pivot table showing the mutated position, its effect and the alternate fraction, representing how many reads contain the mutation (Tab 1). The effects of each mutation are automatically reported in the effect column (EFF) alongside the called variants in our pipeline and include the effect itself, its severity on the affected gene, the changes in the genome, the resulting protein and finally the affected gene. This analysis allows easy comparison of different strains to identify specific mutations, while preserving all quality in a separate table. The filtered and mapped reads are provided for manual inspection of called mutations or for controlling the segregation of intended changes, along with a detailed quality report that can be opened in a browser. Combining multiple sequencing datasets into collections on the Galaxy server allows effective parallel data analysis. The average runtime, excluding wait time for serving capacity, was 55 minutes for Illumina data, 18 minutes for ONT data, and 48 minutes for combined data analysis, making our pipelines a valid tool for routine whole-genome analysis of knock-out and wild-type strains.

**Fig 1:**
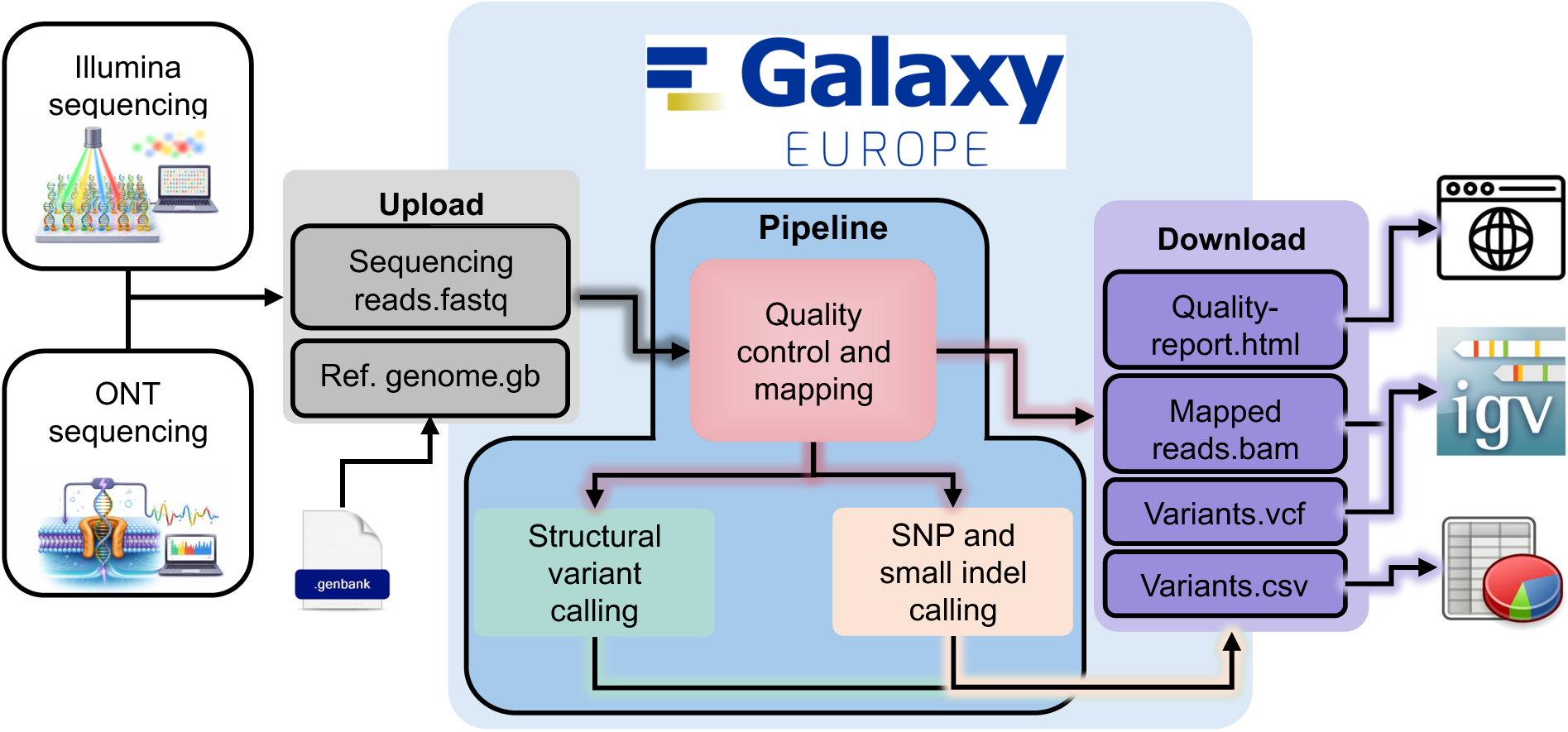
Schematic overview of an easy-to-use galaxy-based pipeline. Schematic overview of the workflows designed and validated here. Each pipeline can be used by simply uploading sequencing data in fastq or fastq.gz format, along with a reference genome, to the European Galaxy server. Starting the pipeline will automatically process the data by performing quality control and mapping, before using different callers for SNP and small indel or structural variant detection. Called variants are filtered and reported as .vcf and .csv files, which can be downloaded along with a detailed quality report and the mapped reads.

**Tab 1:**
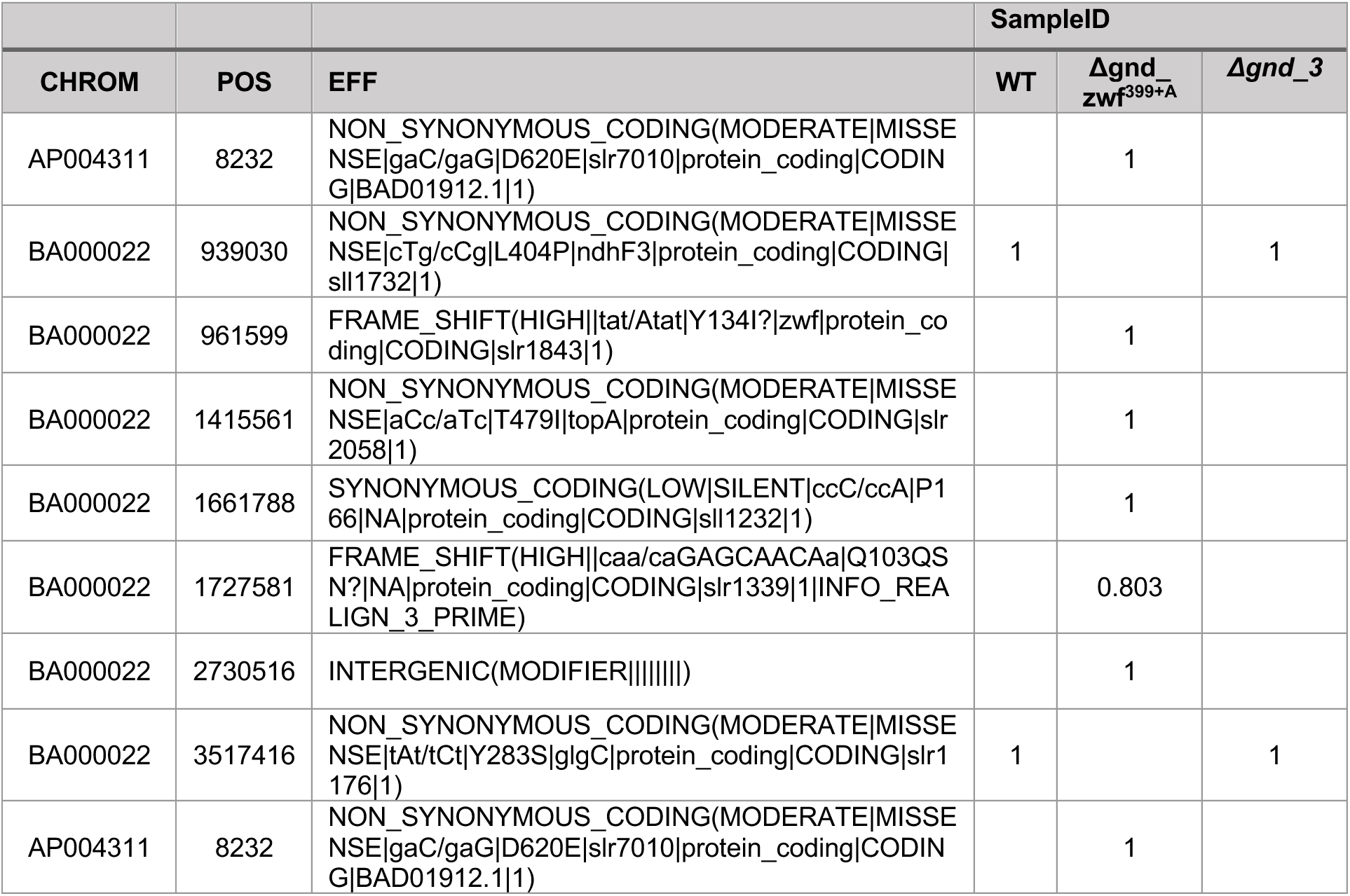
Fixed genetic differences found by Illumina sequencing between wild-type and Δgnd_zwf^399+A^. Mutations were called using Illumina sequencing and the corresponding pipeline. The resulting.csv files were restructured using the provided Excel macro. Only fixed mutations (alternative fraction >0.8) that differ between the two strains are shown. The presented numbers represent the fraction of reads containing the called mutation. Effect collum (EFF) shows the effect on genetic elements on the reference genome (based on SnpEff). Predicted effects are: Effect (effect impact | functional class | codon change | amino acid change| gene name | transcript BioType | gene coding | transcript ID | exon)

### Validation of short and ultra-long read sequencing for variant detection in simulated reads and different *Synechocystis* wild-type strains

We sequenced two non-motile (GT-T) and one motile *Synechocystis* wild-type (PCC-M) substrains previously obtained from labs in Stockholm, Sweden; Uppsala, Sweden; and Freiburg, Germany, using Illumina and ONT sequencing. We then created synthetic genomes containing different types of mutations: Single Nucleotide Polymorphism (SNPs), small insertions and deletions up to 10 bp in size (indels), larger insertions and deletions (up to 2000 bp in size) and complicated structural variants like relocations and duplications (for details see Methods section). Based on these mutated genomes and error profiles derived from the wild-type sequencing experiments, we simulated Illumina and ONT reads with varying coverages. Additionally, we simulated non-segregated variations by mixing simulated reads from the modified genomes with those from the reference genome. Because we used simulated reads, we knew all inserted mutations and used them to calculate the fraction of true positives among all called mutations (precision) and the fraction of true positives that were retrieved (recall or sensitivity) (S11 Figure). We combined by calculating the harmonic mean of the two, the F1-score, as a quality score for each technique (Fig 2). This analysis revealed that in simulated ONT sequencing data, our pipeline struggled with small mutations such as SNPs and indels, especially when the coverage was low or the mutation was not fully segregated. Simple and complex structural variants, however, were detected very reliably even at low coverage or in mixed populations, provided they accounted for more than 15% of the reads (Fig 2A, C). With simulated Illumina data, the pipeline detected SNPs, short indels, and simple structural variations even at low coverage and in non-segregated populations, but struggled with complex structural variations (Fig 2B, D). Based on these results, filters in each pipeline were adjusted, and an additional workflow was created that combines the previous two. In this hybrid version, ONT data are used to detect structural variations, while Illumina data are used to detect smaller changes, such as SNPs and indels (S10 Figure).

**Fig 2:**
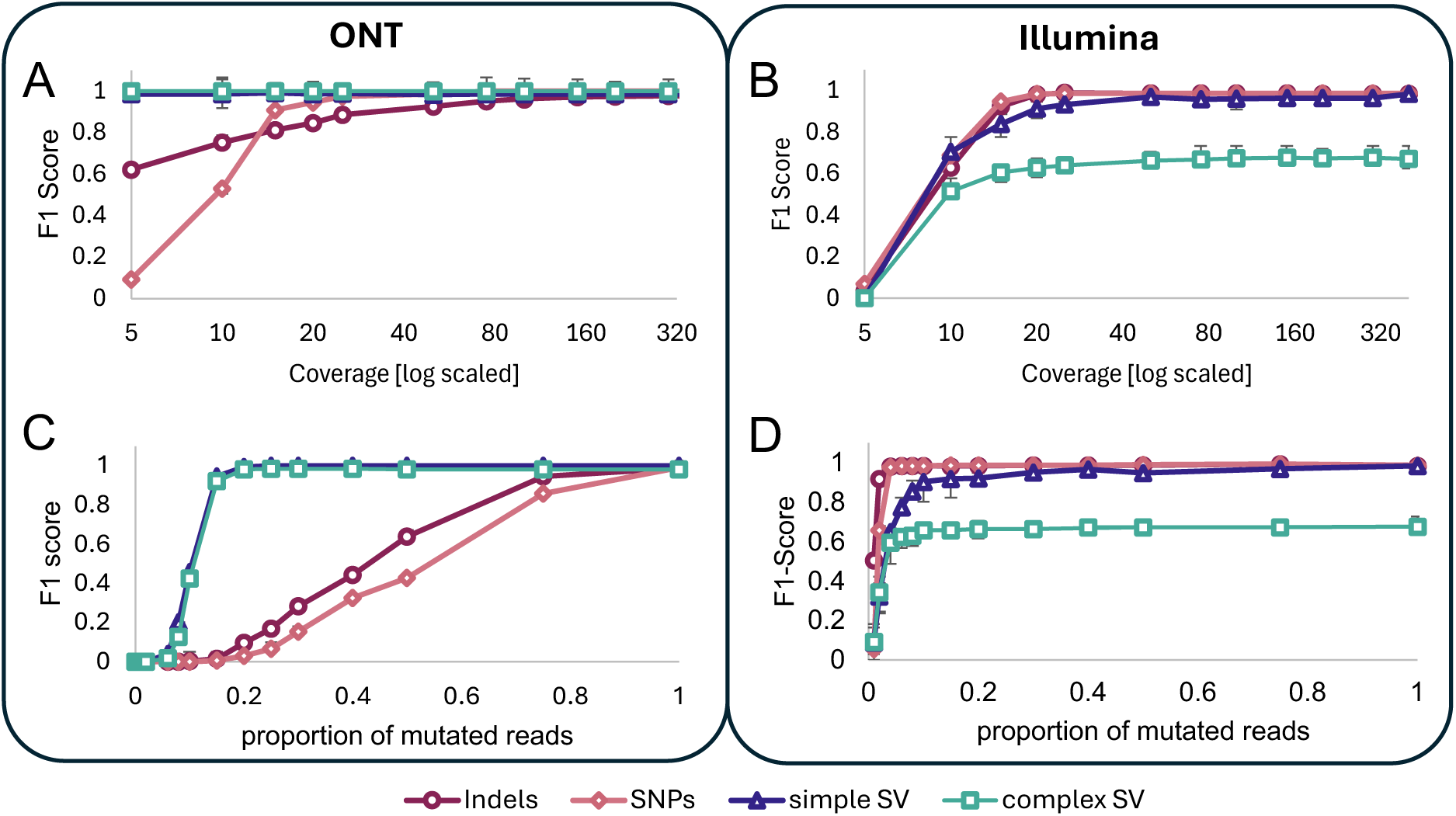
Validation of variant call pipeline for short and long read sequencing data. Variant call pipelines were tested using simulated reads from intentionally altered genomes (see the methods section for details). A) F1-score of variants called from simulated long ONT (Oxford Nanopore Technology) reads with different coverage. B) F1-score of variants called from simulated Illumina reads with different coverage. C) F1-score of variants called from simulated ONT sequencing reads with changing ratios of alternative reads representing non-segregated mutants (combined coverage 225x). (D) F1-score of variants called from simulated Illumina reads with changing ratios of alternative reads representing non-segregated mutants (combined coverage 500x).

Finally, we analyzed the wild-type sequencing data using our pipeline (Fig 3E). Overall, 26 distinct structural variants were identified across all strains; 10 of these (44%) were detected by both techniques, while 12 (52%) were found only in ONT data (Fig 3A). Additionally, many small mutations adjacent to structural variants on extrachromosomal plasmids were called, indicating severe differences from the reference genome at these loci (S6 Table). Apart from these regions, additional small mutations were identified and considered fixed when more than 80% of the reads showed a change at that position. Most of these 55 were identified by both techniques (93%). Additional changes that were not fixed across the population were detected more reliably by Illumina sequencing. Two of the substrains sequenced here (Freiburg and Uppsala) were re-sequenced in other studies before (11,26). Most reported changes were also detected here in both Illumina and ONT sequencing data (Fig 3B, C). However, in both strains, additional fixed mutations were detected here, which have not been reported before. Comparing the three sequenced strains based on structural variants, fixed and non-fixed mutations revealed high similarity between the two non-motile strains, Stockholm and Uppsala. No fixed mutation was detected that was present only in one of these strains, but several non-fixed candidates could be identified. In contrast, several (9) structural variants and many fixed, smaller mutations (21) were found only in the motile strain from Freiburg (S6 Table).

**Fig 3:**
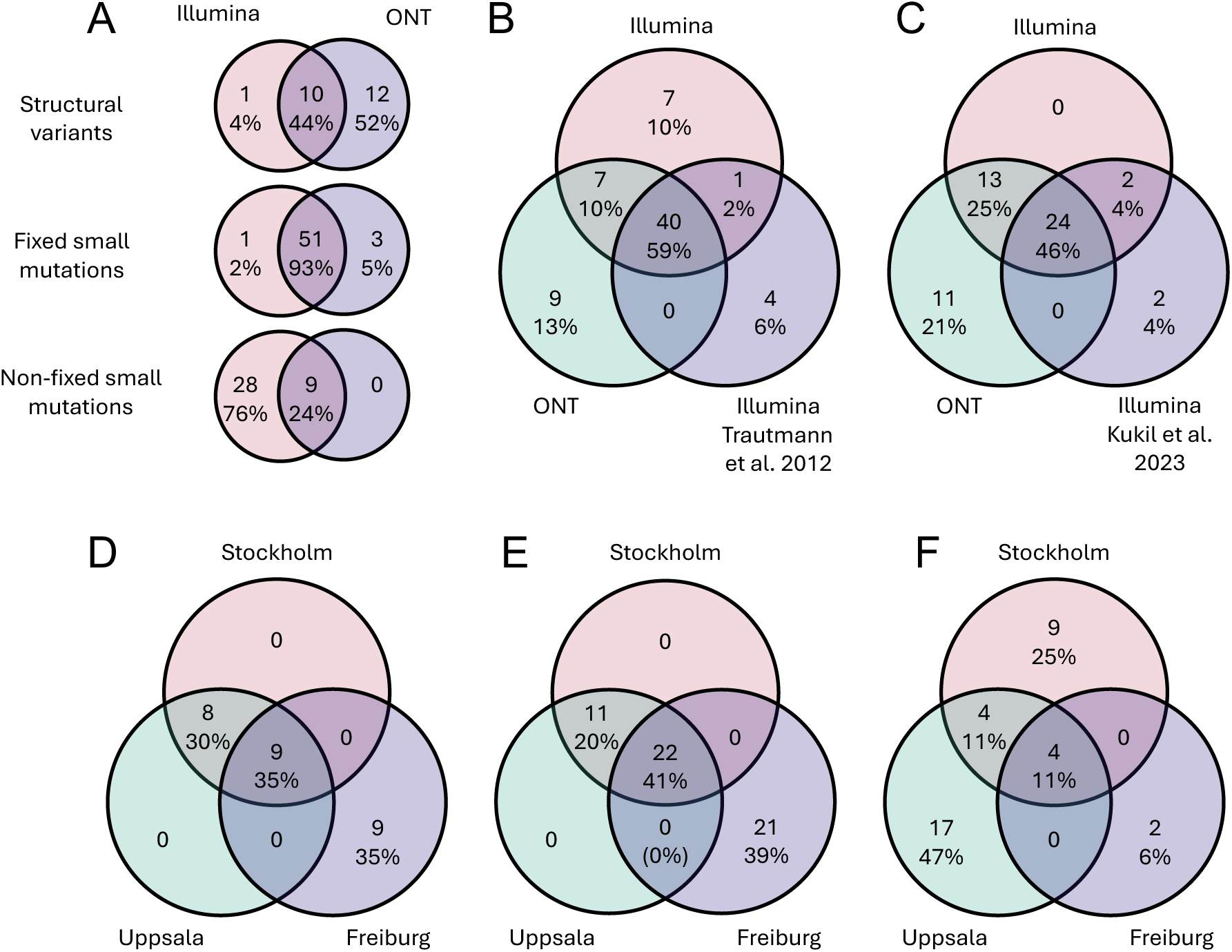
Genetic variations found in different *Synechocystis* wild-type strains using Illumina and Oxford nanopore technology sequencing. Three different wild-type strains from Labs in Stockholm (Sweden), Uppsala (Sweden), and Freiburg (Germany) were sequenced using the Illumina and Oxford Nanopore Technologies (ONT) platforms. Genetic variants were called using different pipelines (see the methods section for details), and the results were compared. A) Venn Diagram of unique structural variants, fixed small mutations (alternative fraction >0.8, length <10bp), and non-fixed small mutations (alternative fraction <0.8, length <10bp) found using Illumina or ONT. B) Comparison of fixed variations (structural variants and small mutations) in the Freiburg strain identified using Illumina sequencing, ONT sequencing, with those published in Trautmann et al. 2012 for this wildtype strain. C) Comparison of fixed variations (structural variants and small mutations) in the Uppsala strain identified using Illumina sequencing, ONT sequencing, with those published in Kukil et al. 2023 for this wildtype strain. D-F) Comparison of the variations found in different wild-type strains for D) structural variants, E) fixed small mutations, and F) non-fixed small mutations.

### Case study of 6-phosphogluconate dehydrogenase mutants reveals the strengths and limitations of different complementation strategies and the impact of secondary mutations

To demonstrate how WGS and different complementation strategies can validate a knock-out phenotype, we compared two strains. One was the previously published *gnd* deletion strain (21), which has recently been renamed to Δgnd_zwf^399+A^ due to a secondary mutation in another key enzyme of the OPP pathway (23). The other set of mutants was generated by re-creating the original plasmid and transforming it into the same wild-type strain (Fig 4A). Six independent colonies were picked and restreaked several times to ensure segregation, which was controlled via PCR (S9 File). After that, three segregated strains *(Δgnd_3_1*, *Δgnd_3_2* and *Δgnd_3_3*, hereafter referred to as *Δgnd_3*) were selected, and their growth under heterotrophic conditions was compared with that of the previous mutant and the wild-type. *Δgnd_zwf^399+A^* grew worse than the wild-type while *Δgnd_3* showed complete growth inhibition (24) (Fig 4B). We complemented both mutants by reintroducing *gnd* at its native locus, which restored heterotrophic growth only in *Δgnd_3*::*gnd,* as expected (Fig 4B, C). We performed Illumina sequencing of both knock-out strains and the corresponding wild-type. Analysis of the data using our pipeline identified six fixed genetic differences unique to gnd_zwf^399+A^ and two mutations only found in the wild-type (Tab 1). The changes found in gnd_zwf^399+A^ were one intergenic mutation, three non-synonymous amino acid changes and 2 frame shifts, including the known mutation in *zwf* (23). The two missense mutations identified in the current wild-type strain were also found in *Δgnd_3*, but no additional changes were identified (S2 Table). Enzyme activity measurements confirmed that only strains containing the zwf mutation, *Δgnd_zwf^399+A^* and *Δgnd::gnd_zwf^399+A^*, had no detectable Zwf activity (4D). To assess whether the increased copy number of plasmid-based expression influences knock-out phenotype rescue, we also reintroduced *gnd* into *Δgnd_3* using plasmid-based expression under the native promoter. This *Δgnd_3* P*_gnd_*:*gnd* strain also regained heterotrophic growth; however, it still grew more slowly compared to the wild-type and the strain complemented on a genomic basis Δ*gnd_3*::*gnd* (Fig 4C). Consequently, we tested Gnd activity in all strains to estimate *gnd* expression levels (Fig 4E). Gnd activity differed markedly between the two strategies: genomic reintegration produced wild-type-like Gnd activity, whereas plasmid-based expression more than doubled it, indicating increased expression levels.

**Fig 4:**
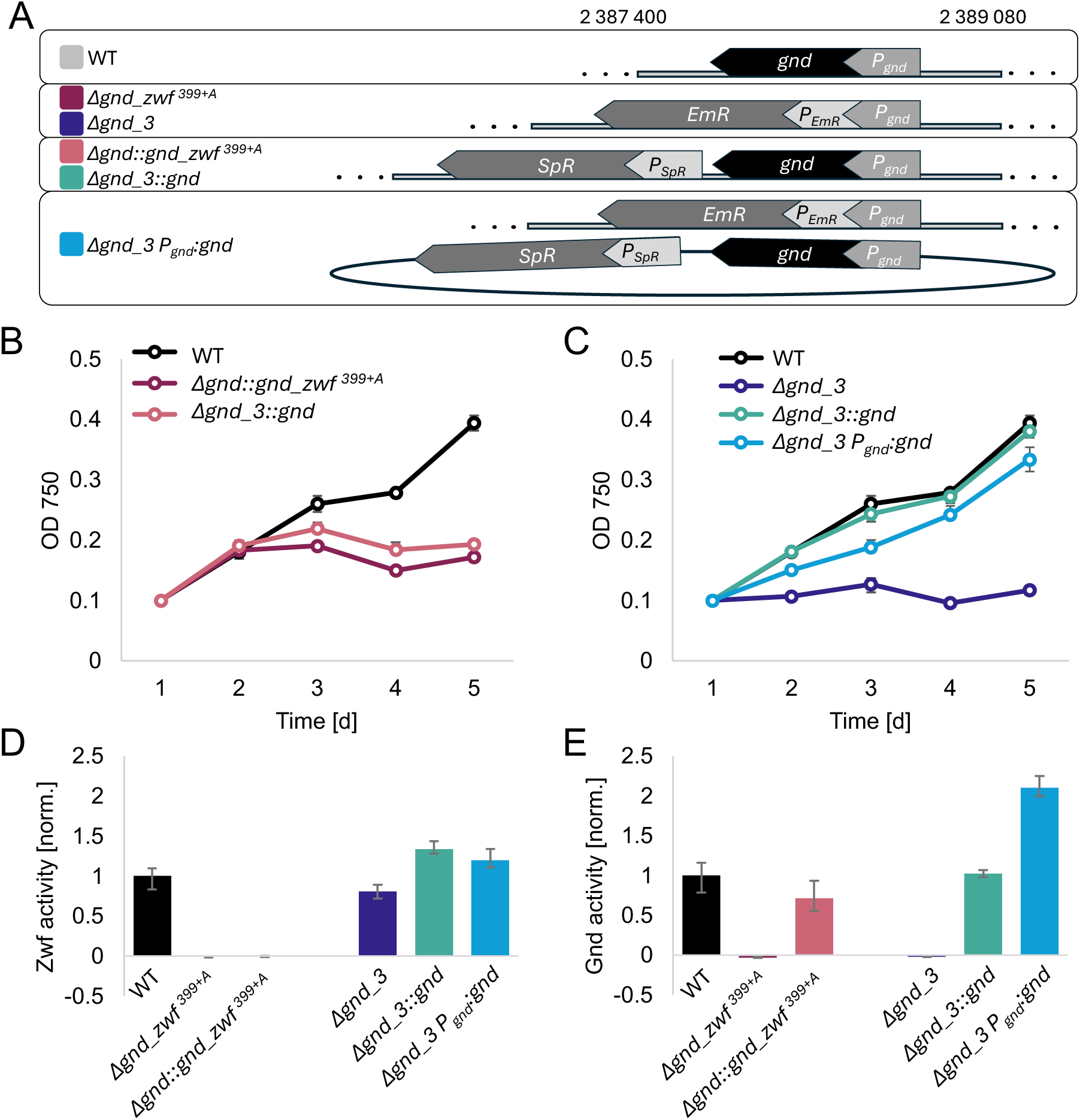
Heterotrophic growth and enzyme activities in different 6-phosphoguconate dehydrogenase (*gnd*) knock-out mutants and complemented strains. **A)** Overview of the strains used. Three dots indicate a continuing genome. **B)** Heterotrophic growth of wild-type *Δgnd_zwf399+A* and *Δgnd_3*) and their corresponding complemented strains were the *gnd* gene was reintroduced in its original locus. **C**) Heterotrophic growth of *Δgnd_3* strain and different complemented strains: *Δgnd_3*::*gnd* reintroduction of the Gnd gene at its original locus, *Δgnd_3* P*_gnd_*:*gnd* reintroduction on a stable plasmid containing the Gnd gene under its original promoter. **D**) Zwf enzyme activity in crude cell extract of the strains used in A and B, normalized to the average WT activity. **E**) Gnd activity in crude cell extract of the strains used in A and B, normalized to the average WT activity.

### The majority of *Synechocystis* studies do not use complemented strains to control a knock-out phenotype

To investigate how common it is to work with knock-out mutants and to control them using their respective complemented mutants in the current *Synechocystis* literature, we conducted a large-scale, AI-based literature analysis. We screened all full-text, open-access research articles from the Europe PMC database that contained *Synechocystis* in the title or abstract using a zero-shot, large-language-model-based classification workflow (N=902; for details and prompts used, see Methods and S8 File). For each paper, we classified: (i) whether the study focused on *Synechocystis*, (ii) whether a new phenotype from a knock-out strain was described, and (iii) whether this phenotype was controlled using a complementation strain (Fig 4E). Validation of the automated approach using a manually classified random sample of 50 papers revealed precision greater than 0.93 and recall greater than 0.89 (S1 Table). This analysis revealed that after removing 178 publications that were not focused on *Synechocystis*, 337 of 740 (46%) studies reported a new phenotype using knock-out mutants, but only 131 of these 337 (39%) used complemented strains as controls (Fig 4). To determine whether this pattern is specific to studies on *Synechocystis* or common in microbiology, we applied the same method and scoring criteria to studies analyzing other model organisms, *Escherichia coli* and *Saccharomyces cerevisiae* (Fig 4B). In these studies, complemented strains were reported in approximately 63% of *E. coli* related publications (3619 of 5715) and in 55% of *S. cerevisiae* studies (2668 of 4876), showing a marked difference in the respective research communities.

### A practical guide to working reliably with *Synechocystis* sp. PCC 6803 knock-out mutants

Finally, we present a step-by-step guide based on our findings, suggesting a workflow for robust knock-out experiments (Fig 5). Detailed considerations for each step are provided (S7 Text), covering key aspects from planning and cloning to mutation generation and experimental design. Focusing on the common strategy of gene replacement via homologous recombination, the guide highlights potential pitfalls and offers practical approaches to minimize them using the complementation and sequencing methods described here. Designing individual studies with these recommendations in mind will improve reproducibility, reduce the risk of misleading phenotypes, and strengthen confidence in the link between observed effects and the intended gene deletion.

**Fig 5:**
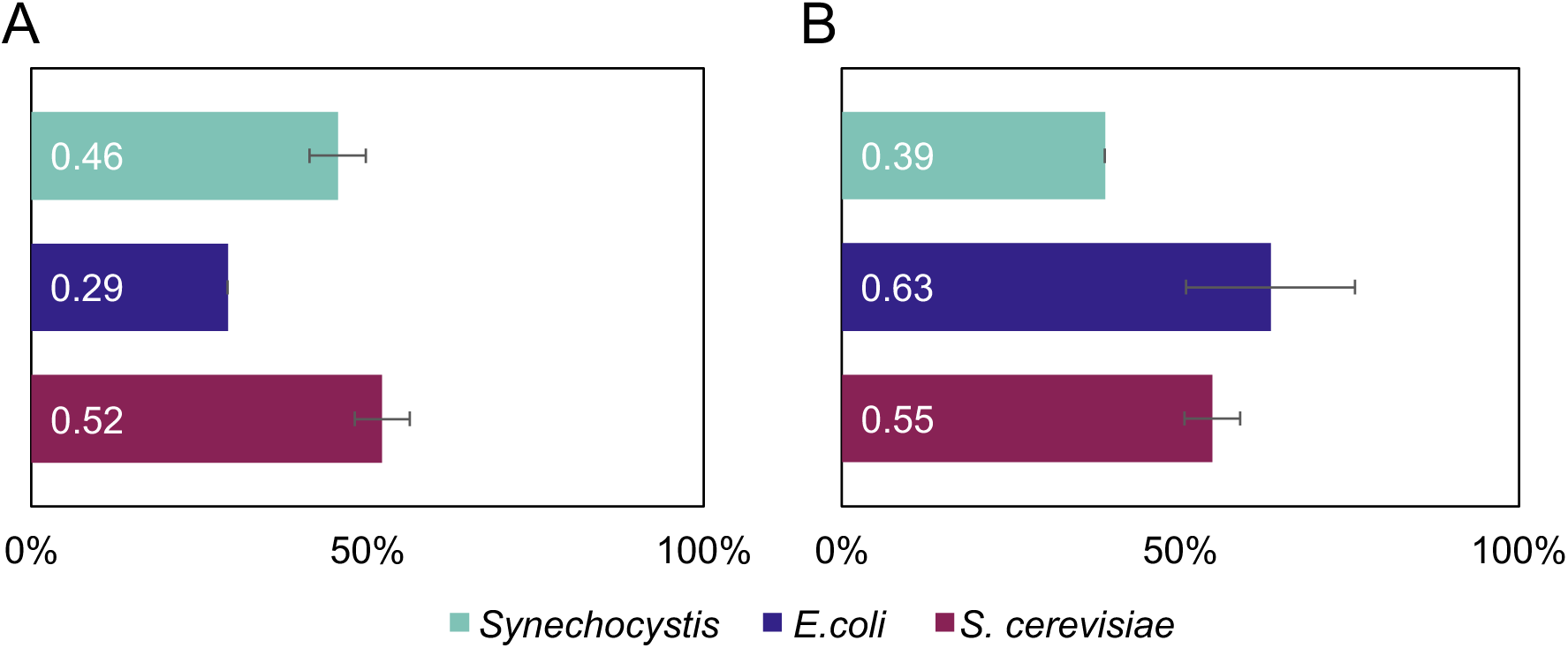
Large language model-based literature analysis about the use of knock-outs and complementation in different model organisms. LLM-based (Mistral-7B-Instruct-v0.3) classification of all full-text, open-access research articles about different model organisms. All publicly available original research articles that mention the model organism in the title or abstract were downloaded from the Europe PMC database (N=901 for “Synechocystis”, 14081 for “cerevisiae”, and 61528 for “coli”). Classified was: whether the study was focused on the model organism (S1_Table), (A) whether a new phenotype from a knock-out was described, and (B) whether this phenotype was controlled using a complementation strain. Error bars represent the average discrepancy relative to 50 random manually classified samples.

**Fig 1:**
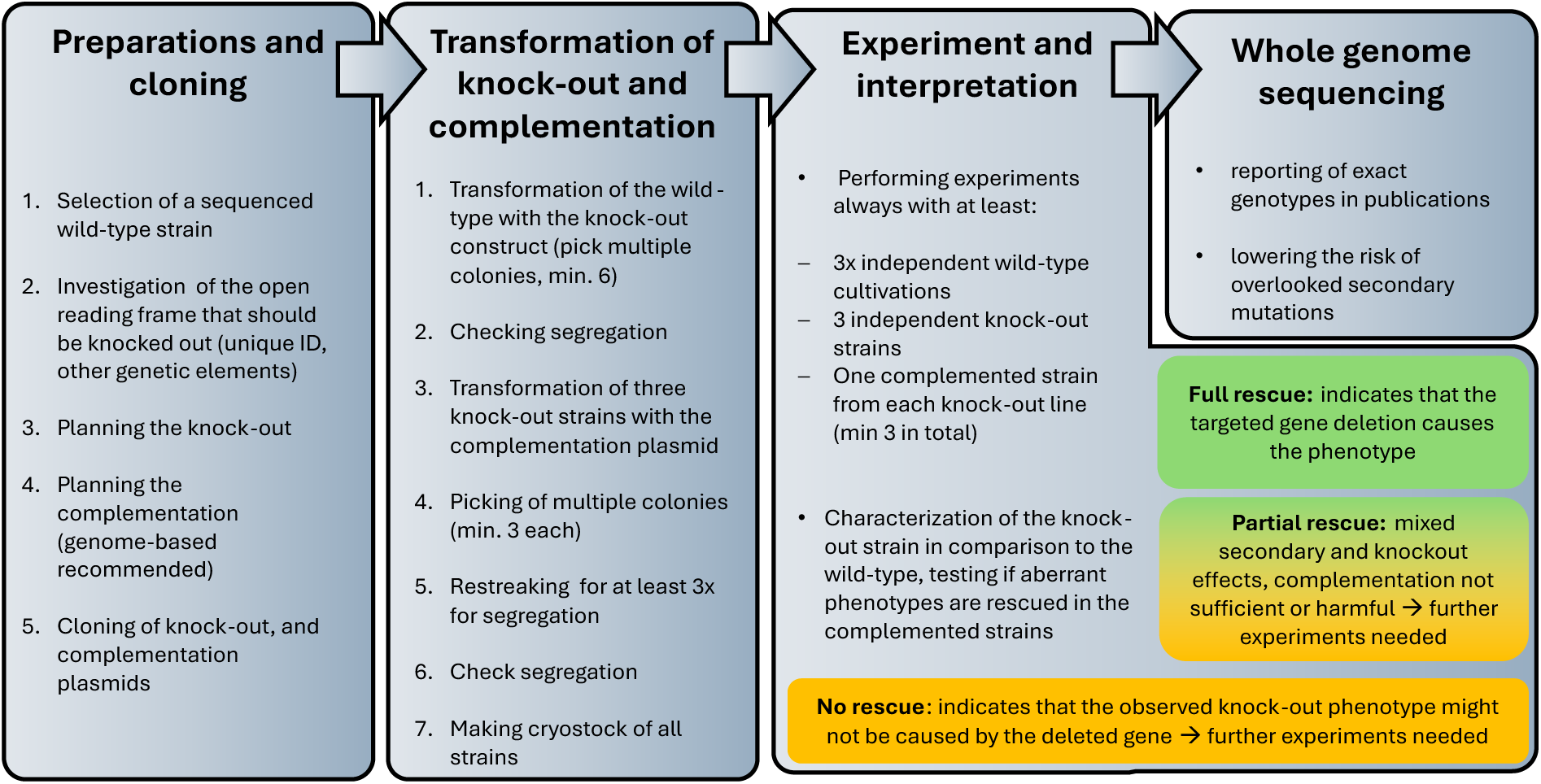
Proposed practical guide to work with knock-out mutants in *Synechocystis* sp. PCC 6803 facilitating robust interpretations. A short, practical guide outlining the main steps for reliable knock-out experiments in *Synechocystis*. Detailed considerations and additional information for many steps highlighted here are available in the supplementary information.

## Discussion

Controlling knock-out mutants using complementation or WGS and reporting the exact wild-type strain used in a study are powerful strategies to make scientific results and publications more reliable and reproducible. Here, we investigated various strategies, highlighted their strengths and weaknesses, and provided tools for streamlining genome sequencing data analysis. By comparing short-read Illumina sequencing to ultra-long read ONT sequencing, we illustrated the known strengths and weaknesses of each technique (15,27). Both techniques and pipelines were employed to detect mutations in three different *Synechocystis* wild-type strains. The distribution of structural variants and fixed and non-fixed mutations identified by Illumina, ONT or both replicated the performance expected from simulated datasets. Comparing these results with previous investigations of the same wild-type strains showed that most known mutations were also found in this study. In addition, several new variants were identified that may be specific to the strains used in this study. However, some especially structural variants were found only by ONT sequencing. While this technique has been used to sequence *Synechocystis* before, no detailed reports on changes compared to the reference genome were made there (Battistuzzi et al., 2024, Synechocystis ATCC 27184)). Given the limitations of prior Illumina sequencing, it is likely that these variants are present in other strains as well, but have not been detected to date (11,26). Comparing the three strains revealed high similarity between the Stockholm and Uppsala strains (GT-T), while clear differences were observed between them and the Freiburg strain (PCC-M). This was expected since Stockholm and Uppsala both belong to the non-motile strain, while the Freiburg strain is motile, and many genetic differences between these lines are known (10,11). While no fixed genetic change was found between the two non-motile strains, the non-segregated mutations may reveal hidden variations either within the population or across different copies of the genome in each cell, which could become strain-specific mutations in the future. While most genetic deviations can be reliably detected using one of the techniques shown here, it cannot be ruled out that rare mutations or those in hard-to-sequence regions are still overlooked. In addition, mutations identified through WGS should always be examined by reviewing the mapped data and, if important to the hypothesis, verified using an independent method.

Technical advances have significantly reduced the cost of sequencing, making it accessible to a wider community. The pipelines provided here, together with the cost-efficient and reliable sequencing techniques available, may facilitate the sequencing of wild-type and mutant strains to obtain additional information on secondary mutations, genetic background, segregation, and hidden variance in the genome. Publishing this information alongside the experimental data generated using the strains will make experiments more reproducible across the scientific community.

We used a case study of two independent *Δgnd* strains to demonstrate how complementation can distinguish true knock-out effects from phenotypes caused by secondary mutations or polar effects. The failed rescue of *Δgnd_zwf^399+A^* indicated that its phenotype relative to that of the wild-type was not only due to the *gnd* deletion but also to secondary mutations. While further experiments are needed to determine which mutation prevented the rescue, the reported effect can indicate the severity of the genetic change (Tab 1). The effect of missense mutations or ones in intergenic regions can usually only be revealed by detailed investigation or additional experiments. The effect of frameshifts, however, is more severe, especially if the mutation does not occur at the end of the open reading frame. The two frame shifts affected *zwf* and slr1339, a gene coding for a small protein of unknown function. Because Zwf is the only enzyme in *Synechocystis* that produces the substrate for *Gnd*, this mutation most likely prevented complementation-mediated rescue (23). The growth phenotype of *Δgnd_3* under heterotrophic conditions was more pronounced than that of *Δgnd_zwf^399+A^*, indicating a distinct phenotype in the *gnd* deletion mutant with an intact *zwf*. This difference highlights the importance of complemented strains as controls, even if the observed knock-out phenotype was expected based on the literature or previous experiments. The slight growth defect observed in *Δgnd_3* P*_gnd_*:*gnd* could indicate that increased Gnd activity from plasmid-based expression affects the cells, revealing a fundamental disadvantage of this strategy. The copy number of RSF1010-based plasmids, such as the one used here, is reported to be 1-3 per chromosome, which is well in line with the measured increase in Gnd activity by a factor of two (29,30). The absence of additional mutations in the independently generated *Δgnd_3* strain relative to its wild-type underscores the importance of working with multiple independent lines, while mutations found in both *Δgnd_3* and the corresponding wild-type indicate lab-specific variations (S2 Table). Even if independent strains would have been enough to prevent misinterpretation in this case, multiple studies reported strains that accumulated mutations in similar regions under high selection pressure, making the use of independent lines an insufficient control and the use of complemented strains essential (8,9).

The AI-based literature analysis revealed that such controls are not currently used in the majority of knock-out studies. While our manual validation of a 50-paper subset indicated high precision and recall, this sample is relatively small (approximately 5.5% of the Synechocystis dataset). Furthermore, large language models can occasionally misinterpret unconventional or highly complex methodological descriptions, particularly if complementation strategies are not explicitly termed as such. Consequently, the exact percentages reported here should be interpreted as a robust statistical trend rather than an absolute metric. Nevertheless, the large sample size of 901 articles provides strong evidence that a significant portion of the community waives these controls. A possible explanation is not a lack of awareness but rather a combination of practical barriers. Due to its polyploidy and slow growth rate compared to other model microorganisms, it usually takes several weeks to generate a single knock-out mutant, which is doubled for complementation (Chauvat et al., 1989). However, skipping these controls can have severe consequences, as shown by investigations regarding the proposed glycolytic Entner-Doudoroff (ED) pathway in *Synechocystis* (21,23). The overlooked secondary mutation in gnd_zwf^399+A^ led to the conclusion that this pathway exists and carries at least a minor flux, despite conflicting evidence (24,31). Recent investigation revealed that both plants and cyanobacteria possess a promiscuous ED aldolase but no ED pathway (23,32,33). Implementing these controls could have prevented this misinterpretation, redirected research efforts earlier, and saved substantial time and resources across multiple laboratories over several years.

To reduce the effort required to implement these controls, we provide tools and guidelines for conducting knock-out experiments. This guide is intended as an easy starting point for the workflow that allows customization for the designated purpose; detailed considerations on how to adjust for each experiment are presented (S7 Fig). Overall, the topic addressed in this paper has apparently been somewhat neglected in parts of the *Synechocystis* community, including our labs. However, with the emergence of novel techniques and their changing availability as costs and efforts decrease, established workflows must be constantly reevaluated and adapted. We strive to improve the robustness and reproducibility of future projects in our lab and others by outlining the necessary steps and providing the tools developed in this study.

## Materials and methods

### Cultivation conditions for enzyme activity assays and growth curves

Strains were maintained on BG11 plates at 28 °C, containing antibiotics if needed, and under constant illumination (50–100 µmol m^−2^ s^−1^).

Pre-cultures for growth experiments were grown autotrophically, first in baffled shake flasks for at least 3 d (in 50 ml BG11). For mutants, antibiotics were used in the pre-cultures, but not in growth experiments, unless noted otherwise. For growth experiments, *Synechocystis* sp. PCC 6803 WT (GT strain obtained initially from the McIntosh lab, Michigan State University USA) and mutants were grown in 200 ml BG11 medium (pH 8) at 28 °C in glass tubes and equally aerated with filter-sterilized air, as described previously (34). Cultures were inoculated at an OD_750_ of around 0.1 to 0.15 and illuminated from two sides with constant light of 50 µmol m^−2^ s^−1^. Heterotrophic cultures were kept in darkness, except for daily illumination of around 10 min, and 10 mM Glucose was added at the beginning of the experiment. To monitor growth, samples were taken from the tube, and OD_750_ was measured daily using a 96-well plate and a plate reader (Infinite M Nano+, Tecan Group AG). In each experiment, three biological replicates were used.

The following antibiotic concentrations were used: kanamycin 50 µg ml^−1^, spectinomycin 20 µg ml^−1^, gentamycin 10 µg ml^−1^ in agar plates, 2.5 to 10 µg ml^−1^ in liquid culture.

### Cloning of the knock-out plasmids

Plasmids used to create the new *Δgnd_3* strain were obtained by first creating a *gnd* knock-out plasmid. Therefore, the genomic region ∼230 bp upstream and downstream of the *gnd* locus was amplified and cloned into a pBluescript plasmid cut with SmaI. Because the *Δgnd_zwf399+A* strain was used as a template, the fragment already contained the Erythromycin resistance cassette and the flanking regions needed for homologous recombination.

### Cloning of the complementation plasmids

To complement the knock-out strains, three different plasmids were created. First, a plasmid for reintegrating the *gnd* gene into its original locus was constructed. Therefore, a region ∼250 bp upstream of the *gnd* gene and the *gnd* gene itself, along with a 250 bp downstream region, were amplified. The fragments were combined into a pBluescript plasmid opened with *Kpn*I and *Sac*I using Gibson assembly. An *EcoR*V restriction site, included in the overlapping primer, enabled the insertion of a spectinomycin resistance cassette downstream of the *gnd* gene and upstream of the *gnd* downstream region.

The other two plasmids were based on the RSF1010-based plasmid pSHDY-Prha-mVenus_rhaS (Addgene [a1] plasmid #137662, https://www.addgene.org/137662/, RRID: Addgene_137662; (35)). From this, extraneous *Nde*I restriction sites were eliminated and a spectinomycin resistance was introduced, resulting in pSSRV. From pSSRV, the intermediate plasmid pSSR_C_and_N_6His was generated, in which mVenus was replaced with a synthetic insert encoding both N-terminal and C-terminal His-tags flanked by unique restriction sites, by removing the mVenus insert by cutting with *Nde*I and *Sti*I and replacing the insert with two fragments with overlapping ends. The complementation plasmid was created by cutting this plasmid with *Pst*I and *Hind*III, removing the insert and the rhamnose inducible promoter and the His-tags, and replacing it with a fragment containing the *gnd* gene and a 130bp upstream region presumably containing the *gnd* promoter.

### DNA extraction and sequencing: Illumina

Cells for Genomic DNA for Illumina sequencing was isolated from *Synechocystis* using a phenol-chloroform method adapted from Makowka et al. (2020). For this, 50 mL liquid culture was centrifuged (8,000 g, 10 min) and the cells were resuspended in 500 µL sterile TE buffer (10 mM Tris and 1 mM EDTA, pH 7.4). Cells were broken by the addition of 10 µL 10 % (w/v) SDS, 500 µL phenol-chloroform-isoamylalcohol solution (25:24:1 (v/v/v)), 500 µL glass beads (0.17-0.18 mm diameter), and by vortexing for 3 min at 4°C. The lysate was then centrifuged (2 min, 10,000 xg, RT) and the upper phase, containing the nucleic acids, was transferred to a new centrifugation cup. For further purification, 500 µL phenol-chloroform-isoamyl alcohol solution (25:24:1 (v/v/v), 4 °C) was added, and the mixture was centrifuged (2 min, 10,000 xg, RT). The upper phase was transferred to a new cup, mixed with 500 µL chloroform-isoamyl alcohol (24:1 (v/v)), and centrifuged (2 min, 10,000 xg, RT). This step was repeated twice, and the resulting supernatant was transferred to a new cup. A volume of 3 M sodium acetate (pH 5), equivalent to around 10% of the supernatant volume, was added, and the cup was mixed by inverting before adding a volume of absolute ethanol (−20 °C) equivalent to around 250% of the supernatant volume. To precipitate the DNA, the samples were incubated at −20 °C overnight, then centrifuged (15 min, 10,000 xg, 0 °C). The supernatant was discarded, and the pellet was washed by adding 1 mL of 70 % (v/v) ethanol (−20 °C), followed by centrifugation (5 min, 10,000 g, RT). After discarding the supernatant, the pellet was dried at 50°C for around 5 min and dissolved in 20 µL TE buffer. Optionally, the DNA concentration was measured with a spectrophotometer (DS-11 FX, Biozym, Hessisch Oldendorf, Germany). Samples were normalized to 20 ng/µl, and 500 ng were sent in for short read NovaSeq 150 sequencing (Mircrobe-EZ, Genewiz, Germany)

### Whole-genome sequencing data analysis

All software, unless stated otherwise, was used on the EU Galaxy server(25).

### Synthetic genome mutations

All scripts used are available in the supplement. Synthetic mutant genomes were generated using custom Python scripts that insert specified mutations at random sites within the reference genome, while generating Variant Call Format files (.vcf) and tab-separated value files (.tsv) that contain all modified sites and were used as truth states(36). For testing SNPs and small INDELS, 200 sites were inserted with a minimum gap of 3500 bp between them, and 20 % of the inserted mutations were elongations or shortening of a single nucleotide repeat of at least 6 bp. INDEL sizes were equally distributed between −10 bp and 10 bp. Larger insertions and deletions were inserted in a similar way. Sizes were 100 bp, 200 bp, 500 bp, 1000 bp, and 2000 bp for insertions and deletions, each repeated 3 times for a total of 30 mutations. Relocations and duplications were performed with equal sizes, also 3 times; sources and targets were chosen with a minimum distance of 3500 bp between them, randomly as before. For relocations, the deletion at the original locus and the insertion at the target were treated as a single mutation, yielding a total of 45 mutations. For each kind of mutation (SNPs, small Indels, simple structural variants, and complex structural variants), five modified genomes were created. Based on the mutated synthetic genomes, synthetic reads were simulated for the Illumina data using ART Illumina (see the supplement for specific tool settings) (37). Different ratios of reads from synthetically mutated genomes and reference ones with a combined coverage of 500x were used to simulate non-fixed mutations. The ONT reads were generated using Nanosim on a local machine with a custom error profile from reads (38). As Nanosim can only operate in’ circular’ mode when given a single sequence, the template genome was split into its constituent contigs, and the reads were merged at the end of read generation to represent circular multi-chromosome sequencing data. Different coverage was created by subsampling simulated reads. Different ratios of reads from synthetically mutated genomes and reference ones with a combined coverage of 225x were used to simulate non-fixed mutations.

### Illumina variant discovery workflow

Short reads are first filtered and trimmed using fastP (39) before being mapped using BWA-MEM2 (2.3) (see supplement for specific tool settings) (40). For structural variant calling, the BAM is passed on to two callers, DELLY (0.9.1) and Manta (1.6) (21,41). These caller create variant f iles (.vcf) independently from each other and were filtered based on their internal quality filters, read depth and quality before being merged using JasmineSV (1.0.11) and being passed onto SNPeff for effect annotation (42–44). Simultaneously, the BAM file created by BWA-MEM2 (2.3) was passed onto two SNP and INDEL callers, FreeBayes (1.3.10) and LoFreq (2.1.5) (45,46). These callers independently create variant files (.vcf), which are filtered for quality, read depth, mapping quality, and strand bias before being merged and passed to SNPeff for effect annotation (42)

### ONT variant discovery workflow

Long reads are first filtered for quality and length using Fastplong before being mapped using Minimap2 (2.28) (see the supplement for specific tool settings) (39,47). For structural variant calling, the BAM file is passed to two callers, namely Sniffles (2.5.2) and cuteSV (2.1.3) (48,49). These caller create variant files (.vcf) independently from each other and were filtered based on quality, internal filters, variant length and reads supporting it filters before having their insertion site and size was refined by IRIS (1.0.5) and subsequently being merged using JasmineSV (1.0.11), to account for the possibility of an caller calling no variants, a pseudo variant was appended to each file and filtered again after merging (44). After merging, the VCF is sorted again and passed on to SNPeff (5.2) for variant effect annotation (43). At the same time, the BAM file created by Minimap2 (2.28) was passed to Clair3 (1.0.10) to call indels and SNPs (50). Its output was again filtered, sorted and additionally normalised and left aligned before the effects were again annotated by SNPeff (5.2) (36).

### Joint Illumina and ONT variant discovery workflow

The joint variant discovery workflow utilized the strengths of both sequencing methods. Large structural variants called by Sniffles (2.5.2) and cuteSV (2.1.3) using ONT reads, SNPs and Indels being called by FreeBayes (1.3.10) and LoFreq (2.1.5) using Illumina reads (45,46,48,49).

### Workflow benchmarking

The outputs of the variant discovery workflows were compared with the true table of synthetic genome mutations at the 5’ site. True positives were counted if the called positions were within a 10 bp window of the true one for SNPs and small INDELS, and a 100 bp window for structural variants. False positives were counted if the called positions were not in the start or end set of the true table, using the same buffer as before.

### DNA extraction and sequencing: ONT

ONT sequencing was performed for three *Synechocystis* WT substrains. The Stockholm strain, generously provided by Paul Hudson, is a non-motile glucose-tolerant substrain (GT-T). The Uppsala strain, a glucose-tolerant, non-motile *Synechocystis* substrain, was kindly gifted by Pia Lindberg (GT-T). The motile PCC-M, strain from Freiburg, which was kindly sent years ago by Annegret Wilde. Cryopreserved *Synechocystis* WT strains were streaked on 1% agar plates with BG11 and incubated in a Multitron HT Incubator (Infors, Switzerland) (30 °C, 150 rpm shaking, 1 % CO2, 75 % humidity, 32 µmol(photons)·m-2 ·s-1. 20 ml BG11 in 100 ml Erlenmeyer flasks were inoculated and cultures were incubated under the same conditions and shaking at 150 rpm until an OD750 nm of ∼7-8. Cells from 2 ml culture were harvested by centrifugation for 15 min at 10,000 rcf, resuspended in 1ml 0.1M EDTA (pH 8), and incubated for 3 ½ h at 300 rpm and 24°C to remove EPS. After centrifugation at 10,000 rcf for 10 min, the pellets were either washed once with 1 ml MiliQ (Freiburg strain) or directly stored at −20°C (Uppsala and Stockholm strains) for at least one night. Cells were resuspended in 400 µl XS buffer (1% potassium ethylxanthogenate, 800 mM NH_4_OAc, 100 mM Tris-HCl pH 7.4, 20mM EDTA, pH8.0, 1% SDS) and incubated for ∼2h at 65°C, with three freeze-thaw cycles in liquid nitrogen. For DNA extraction, 1 volume of phenol:chloroform: isoamyl alcohol (25:24:1) was added and mixed gently by inversion. After centrifugation at 10,000 rcf and 20°C for 10 min, the aqueous phase was retrieved, treated with 5µl RNAseA (4.5U in 10 mM Tris pH8.5, 15mM NaCl) and subsequently incubated at 37°C and RT overnight. Phenol:chloroform:isoamylalcohol extraction were repeated four times before adding 1 volume of chloroform or chloroform:isoamylalcohol. Samples were gently inverted and centrifuged at 10,000 rcf and 20°C for 10 min, and the aqueous phase was re-isolated twice with chloroform:isoamylalcohol. DNA in the aquesous phase was precipitated with 0.6 ml ice-cold EtOH and 0.2M NaCl (final concentration) at −20°C for at least 1 night. DNA was pelleted by centrifugation at 10,000 rcf for 30 min, washed twice with 0.4 ml 70% EtOH, air-dried, and resuspended in 20 µl EB buffer (Tris-HCl) by subsequent incubation at 50°C and RT overnight.

DNA samples were further processed by the Genomics and Transcriptomics Laboratory (GTL) at HHU. The DNA concentration was measured using the Qubit dsDNA HS assay. Sample quality was assessed photometrically using a NanoDropOne. The 260/280 and 260/230 ratios were both ≥1.92. DNA integrity (fragment length) was controlled via capillary gel electrophoresis using the DNF-464 kit and a Fragment Analyzer 5300. The mean fragment size ranged between 24000-35000 bp and all samples had an integrity QC value ≥9.9. The ONT sequencing was performed by GTL on a PromethION using an R10.4.1 nanopore and Kit 14 adapter barcoding, and basecalled with the Highaccuracy model v4.3.0 (400 bps).

### Enzyme activity assays

Glucose-6-P dehydrogenase and 6-P-gluconate dehydrogenase activity was measured by first harvesting 50 ml of cells from the exponential growth phase cultivated in Kniese tubes (as described above) at an OD750 ∼1. Cells were resuspended in 500 µl Tris-HCl 0.1 M pH 7.4 and lysed by vortexing (6 cycles, 10 s max speed) with 200 µl 0.18 mm glass beads. Samples were centrifuged at 12000 rcf for 1 min to remove unbroken cells and glass beads. The supernatant was transferred to a new cup and centrifuged for 20 min at 12000 rcf and 4°C to sediment the membrane fraction. The soluble fraction was transferred to a new cup. The absorption at 650 nm (phycobilisomes) was used to normalize the samples. 50 µl of the normalized cell extract was mixed with 915 µl of Tris-HCl (0.1 M, pH 7.4) and 10 µl 0.1 M NADP^+^. 195 µl of this mixture was added to a 96-well plate three times, and the absorbance at 340 nm was measured for 3 min at 30°C in a plate reader (Tecan, Switzerland). 5 µl glucose-6-P (for glucose-6-P dehydrogenase assays) or 6-P-gluconate (for 6-P-gluconate dehydrogenase assays) was added using the injection system of the plate reader, and the absorption at 340 nm was measured continuously for 10 minutes. The slope after substrate injection was calculated and compared to estimate enzyme activity.

### Plasmids, primers and strains used

Detailed information about the plasmids, primers, and strains used in this study is available in the supplementary information.

### Large language model-based literature analysis

First, all open-access, original research articles containing “*Synechocystis*” were downloaded from Europe PMC as XML files using its API and converted to a CSV file using a Python script (51). Then the large language model Mistral-7B-Instruct-v0.3 (52) was downloaded and installed on the University of Kassel’s high-performance Linux (Rocky Linux 9). Using a custom Python script title, abstract, results, and methods sections of each paper were combined with a prompt containing several definitions, detailed commands on how to format the answer and three questions: i) Does the study present original experimental data obtained using or about *Synechocystis*? ii) Does the study present original experimental data obtained using a *Synechocystis* knock-out mutant? iii) Does the study include experimental data from a complementation/rescue strain to test whether the phenotype is specifically due to the gene knock-out? The answers were phrased to elicit Boolean responses, reasoning, and evidence for each question, and the results were combined with the input data into a single CSV file using Python. Fifty papers were randomly selected and categorized using the same questions to enable precision and recall calculations. All scripts and prompts are provided (S8 File).

## Data availability

All sequencing data are available at BioProject number: PRJNA1450533 https://dataview.ncbi.nlm.nih.gov/object/PRJNA1450533?reviewer=easj36tflkd8u1bk 3mt1pkgavu

## Galaxy Pipelines

Illumina variant calling: https://usegalaxy.eu/u/niklaskueppers/w/illumina-variant-discovery ONT variant calling: https://usegalaxy.eu/u/niklaskueppers/w/ont-variant-discovery Hybrid variant calling: https://usegalaxy.eu/u/niklaskueppers/w/joint-illumina-ont-variant-discovery

## Acknowledgments

ONT sequencing was performed at the Genomics and Transcriptomics Laboratory at Heinrich-Heine University Düsseldorf. We thank Tobias Lautwein and Karl Köhrer for their support. Computational infrastructure and support were provided by the Center for Information and Media Technology at Heinrich Heine University Düsseldorf. The authors acknowledge the support of the Freiburg Galaxy Team: Bioinformatics, University of Freiburg (Germany). We thank Marc von der Heiden, Aleksandar Dobrota and Andrej Dobrota for initial assistance in method testing and pipeline exploring. Many thanks to Annegret Wilde, Pia Lindberg and Paul Hudson for kindly providing the *Synechocystis* WT strains. We thank Michelle Gehringer for the DNA extraction protocol and her valuable advice.

## Author contributions

MT, RF, MB, FP, IMA and KG conceptualized the study. MT, RF, NK, MB, PK and FP investigated, MT visualized, MT, NK, PK, OP and FP development of methology. OP, AW, EO, IMA and KG supervised. IMA, KG acquired funding. MT wrote the original draft, all authors reviewed and edited the manuscript.

## Declaration of generative AI and AI-assisted technologies in the manuscript preparation process

During the preparation of this work, the author(s) used ChatGPT 5.2 (OpenAI, 2025) to correct Python code and ChatGPT 5.2 (OpenAI, 2025), Grammarly and Paperpal to optimize language. After using this tool/service, the authors reviewed and edited the content as needed and take full responsibility for the published article.

## Supporting Information

S1 Table AI based classification data

S2 Table Table of mutations found in *Δgnd* and corresponding wild-type

S3 File *Synechocystis* PCC 6803 reference genome

S4 Text Manual for pipeline usage

S5 Text Macro for Excel to import CSV results and create a pivot table for comparison

S6 Table Mutations found in Synechocystis wild-types

S7 Text Detailed considerations for creating knock-outs in *Synechocystis*

S8 File Python scripts and prompts

S9 File Primers, plasmids and strains used, validation of segregation via PCR

S10 Figure A more detailed overview of the different pipelines

S11 Figure Recall and precision of ONT and Illumina pipelines

