## Supplementary material for "Easy-to-use whole-genome sequencing workflows and standardized practices to uncover hidden genetic variation in *Synechocystis* PCC 6803 wild-type and knock-out strains": S4 Text

### 1 Supplementary data variant pipelines

#### 1.1 Where to find and run the Workflows specified in the paper

All Workflows can be found on the Galaxy Europe server (`https://usegalaxy.eu/workflows/list_published`) under the following names:

| Available Workflows for Variant discovery | | | |
|---|---|---|---|
| **Workflow name** | **Workflow features** | **Required inputs** | **Workflow output** |
| ONT_variant_discovery | Accurate structure variant detection.<br><br>Detection of high frequency SNPs possible. | Nanopore reads as .fastqsanger.gz format (see Manual below how todo this).<br><br>Reference genome file in .genbank format | VCF file containing all detected variants, separated by structural variants and SNPs and Indels, as-well as a tab separated file and .csv file. |
| Illumina_variant_discovery | Detection of some Structural variants possible.<br><br>High accuracy SNP and Indel calling. | Illumina paired reads (see Manual below how todo this).<br><br>Reference genome file in .genbank format | VCF file containing all detected variants, separated by structural variants and SNPs and Indels, as-well as a tab separated file and .csv file. |
| Joint_Illumina_ONT_variant_discovery | Accurate structure variant detection.<br><br>High accuracy SNP and Indel calling. | Nanopore reads as .fastqsanger.gz format (see Manual below how todo this).<br><br>Illumina paired reads (see Manual below how todo this).<br><br>Reference genome file in .genbank format | VCF file containing all detected variants, separated by structural variants and SNPs and Indels, as-well as a tab separated file and .csv file. |

#### 1.2 Manual - how to use the workflows

##### 1.2.1 Accessing the Workflows

First you will need to go to https://usegalaxy.eu/ and create an account. After that you can navigate to workflows and press on "Public Workflows" (https://usegalaxy.eu/workflows/list_published) there you will find all published workflows, and also ours used in this paper. If you search for ONT_variant_discovery one workflow should appear.

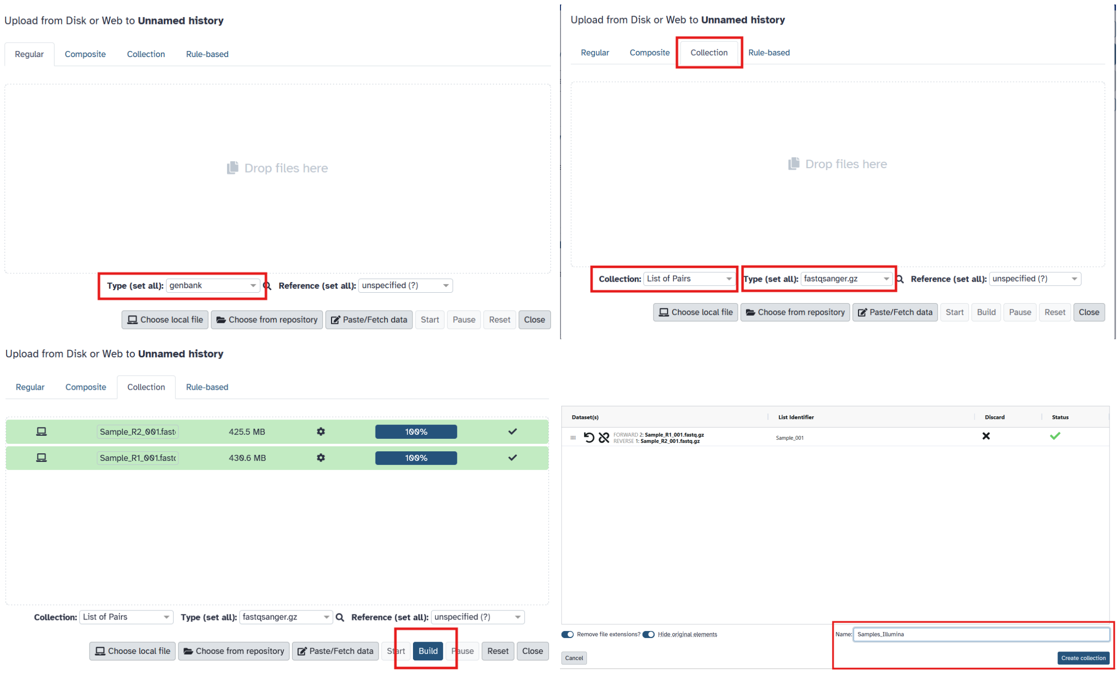


Figure 1: Illumina read upload

##### 1.2.2 Uploading input data

You will need to upload all of your sequencing reads and references to the server. In the case of your .genbank reference you can upload a singular file, just make sure you have set the type to .genbank before uploading. Next step is uploading the paired read data. For that you need to switch to the Collection tab and select type list of pairs, and .fastqsanger. After all samples are uploaded you need to first press 'Build' to actually build the list. After this you can give the List a name and create the dataset After this you can upload your nanopore data, you again need to upload it in a list format and build the dataset.

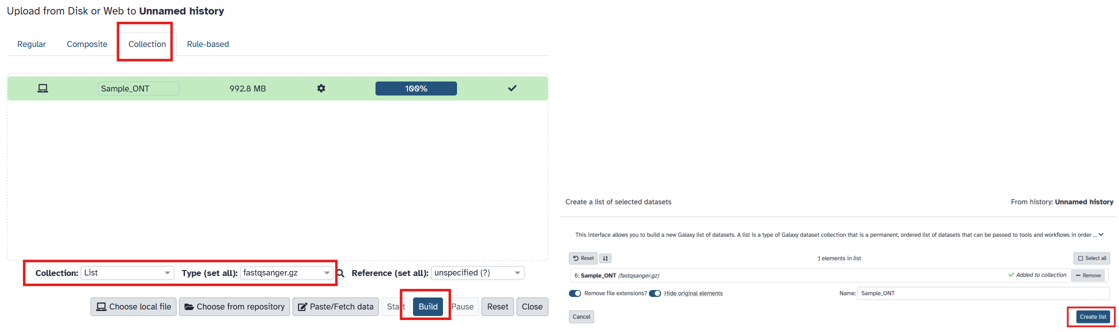


Figure 2: ONT Build

##### 1.2.3 Obtaining output data

Once you have started the workflow and executed it, you will find the invocation history on the left under 'Workflow Invocations', there you can see the current state of the workflow that will also be shown to you after starting. Once the workflow is finished you will find all the workflow outputs in the history you selected to run the workflow in.

#### 1.3 Workflow settings

##### 1.3.1 Variant discovery tool settings

| **Variant discovery pipelines** | | |
|---|---|---|
| **Tool name** | **Tool version** | **Tool settings** |
| SnpEff | 5.2 | Default |
| Fastplong | 0.4.1 | Adapter trimming enabled with defaults, **–trim_front 50**, **phred>=Q10 or userdefined**, **–length_required 500 or userdefined** |
| Minimap2 | 2.28 | **x map-ont** , **-N = 3** |
| QualiMap BamQC | 2.3 | Default |
| CuteSV | 2.1.3 | **Genotyping = Yes**, |
| Sniffles | 2.5.2 | –minsupport 'auto', –max-splits-kb '0.1', –minsvlen '50', –mapq '20', –detect-large-ins '0', –cluster-binsize '100', –cluster-r '2.5', **–minalignmentlength = 150** |
| Bcftools | 1.22 | Dependend on workflow |
| Iris | 1.0.5 | **–rerunracon**,**–also_deletions** |
| JasmineSV | 1.0.11 | **–output_genotypes**, **–allow_intrasample** |
| MultiQC | 1.27 | Default |
| Clair3 | 1.0.10 | **r1041_e82_400bps_hac_v410** |
| Fastp | 0.24.1 | **phred>=Q30**,**minlenght>=150** |
| BWA-MEM2 | 2.3 | Default |
| Filter BAM | 2.5.2 | For SVs Illumina (**Read IS paired**,**Read IS in proper pair**), For SNPs Illumina (**isPrimaryAlignment**, **isMapped**, **isMateMapped**, **mapQuality>=20**) |
| BamLeftAlign | 1.3.10 | Default |
| SortSam | 3.1.1.0 | Coordinate based |
| Delly call | 0.9.1 | **–svtype ALL**, **–map-qual 20**, **–qual-tra 20**, **–mad-cutoff 6**, **–minclip 10**, **–min-clique-size 5**, **–minrefsep 50**, **–maxreadsep 750**, **–geno-qual 3** |

| | | |
|---|---|---|
| Manta | 1.6 | Default: –minCandidateVariantSize '8', –rnaMinCandidateVariantSize '1000', –minEdgeObservations '3', –graphNodeMaxEdgeCount '10', –minCandidateSpanningCount '3', –minScoredVariantSize '0', –minDiploidVariantScore '0', –minPassDiploidVariantScore '0', –minPassDiploidGTScore '0', –minSomaticScore '0', –minPassSomaticScore '0', –enableRemoteReadRetrievalForInsertionsInGermlineCallingModes '1' |
| FreeBayes | 1.3.10 | –min-coverage 50 –skip-coverage 0 –limit-coverage 0 –theta 0.001 –ploidy 1 -K -n 0 –haplotype-length -1 –min-repeat-size 8 –min-repeat-entropy 1 –standard-filters –min-mapping-quality 30 –min-base-quality 0 –min-supporting-allele-qsum 0 –min-supporting-mapping-qsum 0 –read-indel-limit 1000 –min-alternate-fraction 0.05 –min-alternate-qsum 0 –min-alternate-count 5 –min-alternate-total 1 |
| LoFreq | 2.1.5 | –call-indels –min-cov 50 –max-depth 1000000 –min-bq 30 –min-alt-bq 30 –min-mq 30 –max-mq 255 –min-jq 0 –min-alt-jq 0 –def-alt-jq 0 –sig 0.01 |

##### 1.3.2 Variant discovery filter settings

<table>
<tr><th colspan="2">Variant discovery pipelines filters</th></tr>
<tr><th>Filter position</th><th>Filter settings</th></tr>
<tr><td>Bcftools filter vcf after Delly</td><td>(INFO/PRECISE=0) || (INFO/PE < 3 && INFO/SR < 8) || (INFO/PE >= 3 && INFO/MAPQ < 20) || (INFO/SR >= 3 && INFO/SRMAPQ < 30) || (INFO/SVTYPE="INV" && INFO/SR < 10) || (FMT/GQ != "." && FMT/GQ < 20)</td></tr>
<tr><td>Bcftools soft filter vcf after FreeBayes</td><td>INFO/MQM < 30 || INFO/SAP > 30 || INFO/EPP > 30 || INFO/RPP > 30 || INFO/RPPR > 30 || INFO/EPPR > 30</td></tr>
</table>

<table>
<tr><td>Bcftools soft filter vcf after Lowfreq</td><td>(INFO/SB > 20) || (INFO/HRUN >= 8 && INFO/AF < 0.10) || ((INFO/DP4[2] + INFO/DP4[3]) < 5) || (INFO/AF < 0.05)</td></tr>
<tr><td>Bcftools filter vcf after Clair3</td><td>FMT/DP<50 || FMT/AF<0.60 || FORMAT/AD[0:1]<30</td></tr>
<tr><td>Bcftools soft filter vcf after CuteSV</td><td>FILTER!=”PASS” || INFO/RE < 5 || FORMAT/DV < 3 || FORMAT/GQ < 15 || (INFO/SVLEN!=”.” && ABS(INFO/SVLEN) < 50) || (INFO/SVLEN!=”.” && ABS(INFO/SVLEN) > 100000) || INFO/AF < 0.05</td></tr>
<tr><td>Bcftools soft filter vcf after Sniffels</td><td>(FILTER!=”PASS” && FILTER!=”GT”) || INFO/SUPPORT < 10 || INFO/AF < 0.10 || (INFO/SVLEN!=”.” && ABS(INFO/SVLEN) < 50) || (INFO/SVLEN!=”.” && ABS(INFO/SVLEN) > 100000) || (INFO/END!=”.” && ABS(INFO/END-POS) > 100000) || (INFO/STDEV_POS!=”.” && INFO/STDEV_POS > 20) || (INFO/STDEV_LEN!=”.” && INFO/STDEV_LEN > 50)</td></tr>
</table>
