## Supplementary material for "Easy-to-use whole-genome sequencing workflows and standardized practices to uncover hidden genetic variation in *Synechocystis* PCC 6803 wild-type and knock-out strains": S7 Text

### Practical guide to work with Knock-out mutants in Synechocystis- Additional considerations

#### Planning and cloning

While rigorous planning is key in every experiment, we noticed that in apparently simple knock-out experiments, important considerations are often overlooked.

1. Sequence and choose the wild-type strain

As discussed earlier, a wide variety of *Synechocystis* PCC 6803 strains are used and each lab strain can harbor additional mutations. Before starting, it is important to choose the correct wild-type strain, especially when comparing data with previously created strains. Additionally, this wild-type strain should be sequenced (if not done recently) to determine its genetic background. This information should be reported alongside any data generated using this strain.

2. Investigate the open reading frame that should be knocked out

The genomic region to be manipulated must be thoroughly investigated. Besides the GOI, there might be additional elements, such as antisense RNAs, overlapping transcripts, or promoter regions, that would also be influenced. While deleting them cannot always be avoided, it is important to understand these elements and interpret the data accordingly, or implement additional controls. Over the last few years, there have been significant improvements in the annotation of the Synechocystis genome, including the discovery of many non-coding RNAs, small open reading frames, and alternative start sites (Mitschke et al., 2011; Spät et al., 2023; Hadjeras et al., 2026). Therefore, labs must use the most recent versions of the annotated genomes. The annotations provided here (SUPP FILE, based on (Mitschke et al., 2011)), already include experimentally determined transcription start sites and many small RNAs. In addition, the cyanobacteria community is also aware of the problem and is working on a new, well-curated database (Moore et al., 2024).

3. Plan the knock-out

There are different strategies available to knock out a gene in Synechocystis. Knock-out mutants can be generated by replacing the gene of interest with a selection marker, by inserting the marker into the open reading frame via homologous recombination, or by introducing a targeted mutation using a CRISPR-based method (Berla et al., 2013; Cengic et al., 2022). In the first case, which is currently most common, the selection marker can be removed later to create so-called marker-less mutants (Viola et al., 2014). In all cases, the genomic region at that position is affected to varying degrees, which may lead to phenotypes not directly linked to the deleted gene, so-called polar effects (Hutchison et al., 2019). Additional considerations must be made regarding the selection marker. These usually consist of a cassette containing a constitutive promoter, an antibiotic resistance gene, and a terminator. A leaky terminator could influence the expression of downstream genes, again creating an unrelated phenotype. One way to minimize this effect is to first choose a well-characterized terminator for Synechocystis (Kelly et al., 2019) and, if possible, orient the resistance cassette so that it does not face the same direction as the downstream gene. If the GOI is part of a transcriptional unit (an operon), the insertion of a selection marker could influence the transcription of downstream genes by artificially terminating elongation (Nefedova et al., 2003; Mateus et al., 2021). Finally, the selected resistance gene and antibiotic exposure can influence the mutated cells and affect results when comparing the mutant strain to the wild type, even if, in this specific experiment, both cultures were cultivated without antibiotics. Therefore, creating a wild-type-like strain with the same resistance as the mutant by inserting it at a neutral site can be beneficial.

4. Decide on a complementation strategy

As shown in this study, a powerful and important method to separate true phenotypes from secondary effects is to reinduce the deleted gene by creating a complementation strain. There are two main strategies for reintroducing a gene into the Synechocystis knock-out strain. The recombination of the gene back into its original locus on the genome, replacing the knock-out selection marker with another marker, or the transformation with a stable plasmid carrying the gene of interest under the control of either the native or a heterologous promoter. Because many promoter regions are not characterized well in Synechocystis, it is common practice to use up to several hundred base pairs upstream of the gene, assuming it contains all critical elements for native expression. Even with a native promoter, the achieved protein level may vary due to differences in plasmid copy number relative to the genome, as shown in our case study (FIG XX). Because of incorrect annotations of some genes, especially translational start sites, in *Synechocystis*, one could easily truncate a gene by accident during reinduction (Mitschke et al., 2011). Because differences in expression can influence the phenotype or cause new ones, we recommend reinducing the gene at the original locus.

#### Transformation of knock-out and complementation

1. Transform the knock-out construct

Once all plasmids are cloned, the knock-out plasmids can be transformed into the selected wild-type strain. To minimize the risk of a critical secondary mutation, multiple independent strains should be picked after transformation. We recommend picking at least six colonies. While it is highly unlikely that independent strains harbor the same mutation, if not inherited, high selection pressure can lead to the emergence of strains with mutations in similar regions or pathways, so-called suppressor mutants (Kobayashi et al., 2005; Nishijima et al., 2015). Therefore, while increasing the number of independent strains, complementation remains essential to demonstrate that the induced gene deletion directly causes a newly described phenotype. If suppressors occur, having multiple of them allows it to investigate these mutations later, yielding information about the selection pressure in the knock-out mutant.

2. Check segregation via PCR

Later, segregation of the transformed strain is usually confirmed by PCR. Primers flanking the GOI locus are used to amplify the remaining copies of the GOI or the selection marker. However, a simple PCR is not quantitative and will overrepresent fragments favored by the PCR, so fragment size and PCR settings have been chosen accordingly (Walsh et al., 1992).

3. Transform three knock-out strains with the complementation plasmid

The complementation plasmid should be transformed into at least three of the knock-out strains. Again, multiple colonies from each transformation should be picked. To keep the number of mutans manageable, recommend picking six; after segregation is confirmed, keep and cryostock only three, for a total of 12 strains. If all nine complementation strains show similar phenotypes, one from each knock-out strain can be chosen to begin the experiment. If double mutants are created, one should proceed similarly by creating at least one double-knockout from each original knock-out strain to keep the lines as independent as possible.

7. Make cryostock of all strains

After PCR confirms segregation, cryostocks of the generated knock-out and complementation strains should be prepared. Longer cultivation increases the risk of secondary mutations. However, thawing a cryostock can select for the fittest cells, which may favor suppressor mutants, so repeated freeze-thaw cycles should be avoided (Trautmann et al., 2012). While not strictly necessary, given the low cost and effort, one could consider sequencing the strains selected for the experiment to ensure that no unnoticed secondary mutations occur and that the genetic manipulation is fully segregated. When stains are exchanged between labs, resequencing them, as is regularly done with plasmids, can confirm that no errors occurred and ensure that the appropriate wild-type is chosen.

#### Experiment and interpretation

We strongly suggest not starting any experiment without the complete set of wild-type and three independent knock-out lines, and their corresponding complementation strains. The risk of observing an unreliable phenotype in the initial knock-out strains, especially if only one clone is picked, should not be underestimated. Additionally, this usually leads to more experiments investigating this uncontrolled phenotype, which is often more work than creating the complementation strain in the first place. Repeating all the experiments at that stage usually sets the whole project back.

If a knock-out phenotype is observed in the knock-out and is fully rescued by complementation, with no additional phenotypes emerging, this provides strong evidence that the targeted gene deletion directly causes the phenotype.

In contrast, partial rescue is more difficult to interpret. It may indicate that expression of the complemented gene was insufficient to restore native function, or that other factors, such as secondary mutations or polar effects on nearby genomic regions, contribute to the phenotype alongside the deletion. Disentangling these possibilities requires additional, targeted experiments. A failing complementation experiment does not completely falsify the causal connection between the knock-out and the phenotype, it should prevent one from concluding that this connection exists, unless other strong experimental evidence indicates otherwise. Finally, whole-genome sequencing can be used to characterize the investigated mutants in greater detail. Sequencing strains with a phenotype that cannot be complemented can explain this behavior, as in our example, or open new research questions by revealing a mutation in an unexpected gene.
