## Supplementary material for "Easy-to-use whole-genome sequencing workflows and standardized practices to uncover hidden genetic variation in *Synechocystis* PCC 6803 wild-type and knock-out strains": S8 File: Analyze_Paper_LLM_cer.py

```
import os
import re
import pandas as pd
import torch
from transformers import AutoTokenizer, AutoModelForCausalLM
from tqdm import tqdm
import csv

# === CONFIGURATION ===
QUERY_TERM = "cerevisiae"
BASE_DIR = f"./data/{QUERY_TERM}"
RAW_ARTICLES_DIR = f"{BASE_DIR}/raw_articles"

LLM_LABELED_DATA_PATH = f"{BASE_DIR}/llm_batch_labeled_articles.csv"
PARSED_DATA_PATH = f"{BASE_DIR}/parsed_articles_with_metadata.csv"
NUM_ARTICLES_TO_PROCESS = 500000

# === LOCAL MODEL SETUP ===
MODEL_PATH = "/gpfs/home28/fb10-pfph/alphafold/Mistral-7B-Instruct-v0.3"

print("Loading local model and tokenizer...")
tokenizer = AutoTokenizer.from_pretrained(MODEL_PATH, local_files_only=True)
if tokenizer.pad_token is None:
    tokenizer.pad_token = tokenizer.eos_token

model = AutoModelForCausalLM.from_pretrained(MODEL_PATH,torch_dtype=torch.float16,device_map="auto",local_files_only=True)
model.eval()
print("Model loaded successfully.")

# === LOAD PARSED ARTICLE DATA ===
df_parsed = pd.read_csv(PARSED_DATA_PATH, sep=";").fillna("")

# === LOAD PREVIOUSLY LABELED RESULTS (if available) ===
if os.path.exists(LLM_LABELED_DATA_PATH):
    results_df = pd.read_csv(LLM_LABELED_DATA_PATH, sep=";").fillna("")
    labeled_pmcs = set(results_df["pmcid"].tolist())
    print(f"Resuming batch labeling: {len(labeled_pmcs)} already processed.")
else:
    results_df = pd.DataFrame()
    labeled_pmcs = set()
    print("Starting new batch labeling.")

# === FILTER ARTICLES THAT STILL NEED TO BE LABELED ===
remaining_df = df_parsed[~df_parsed["pmcid"].isin(labeled_pmcs)].copy().reset_index(drop=True)

# === PARSE MODEL RESPONSES ===
def parse_answers(response_text):
    pattern = re.compile(
        r"(?:Answers?\s*:?\s*)?"                     
        r"(?:(\d+)\.?\s*)?"                         
        r"(Yes|No)\s*"                               
        r"(?:\n|\r|\s)*Reasoning\s*[:\-]?\s*"  
        r"(.+?)(?=(?:\n(?:\d+\.\s*|Answers?\s*\d*[:\-]?|Answer\s*\d*[:\-]?))|\Z)",
        re.DOTALL | re.IGNORECASE
    )

    matches = pattern.findall(response_text)

    answers = []
    for _, answer, reasoning in matches:
        reasoning_words = reasoning.strip().split()
        reasoning_text = ' '.join(reasoning_words[:50])

        answers.append({
            "answer": answer.strip().capitalize(),
            "reasoning": reasoning_text or "No reasoning extracted."
        })

    while len(answers) < 3:
        answers.append({
            "answer": "Unknown",
            "reasoning": "No reasoning extracted."
        })

    return answers[:3]


# === RUN INFERENCE WITH THE MODEL ===
def ask_model(title, abstract, methods, results):
    prompt = f"""
You are an expert scientific assistant. Carefully read the following sections of a scientific paper:

TITLE:
{title}

ABSTRACT:
{abstract}

METHODS:
{methods}

RESULTS:
{results}

Answer the following three questions. For each question, provide your answer and a short explanation of your reasoning. Use the exact format shown below. Even if the answer is obvious, explain your reasoning clearly.Copy the example format exactly, including the Reasoning section for every question.

Answer: <Yes or No>
Reasoning: <maximum 50 words reasoning for your answer>

Questions:
1. Does the study present original experimental data obtained using {QUERY_TERM}?
2. Does the study present original experimental data and a new phenotype of a {QUERY_TERM} knockout mutant?
3. Does the study explicitly consider alternative explanations for the phenotype of the knockout mutant (e.g., off-target effects, secondary mutations, or experimental artifacts) and use appropriate controls to address them?
"""

    try:
        inputs = tokenizer(prompt, return_tensors="pt", padding=True).to(model.device)

        with torch.no_grad():
            outputs = model.generate(
                **inputs,
                max_new_tokens=300,
                do_sample=False,
                eos_token_id=tokenizer.eos_token_id,
                pad_token_id=tokenizer.pad_token_id,
            )

        responses = tokenizer.batch_decode(outputs, skip_special_tokens=True)
        cleaned_responses = [r.replace(prompt.strip(), "").strip() for r in responses]

        return cleaned_responses

    except Exception as e:
        print(f"Error during model inference: {e}")
        return [""]

# === PREPARE NEW COLUMNS FOR RESULTS ===
print(f"Preparing columns for {min(NUM_ARTICLES_TO_PROCESS, len(remaining_df))} articles.")
for i in range(1, 2):
    remaining_df[f"sample_{i}_raw_response"] = ""
    for q_idx in range(1, 4):
        remaining_df[f"sample_{i}_q{q_idx}_answer"] = ""
        remaining_df[f"sample_{i}_q{q_idx}_reasoning"] = ""

# === PROCESSING LOOP ===
print(f"Starting batch processing for {min(NUM_ARTICLES_TO_PROCESS, len(remaining_df))} articles.")

def clean_text(text):
    if not isinstance(text, str):
        return text
    return ' '.join(text.strip().split())

for index, row in tqdm(remaining_df.head(NUM_ARTICLES_TO_PROCESS).iterrows(),
                       total=min(NUM_ARTICLES_TO_PROCESS, len(remaining_df))):
    pmcid = row["pmcid"]

    if pmcid in labeled_pmcs:
        print(f"Skipping {pmcid}, already processed.")
        continue

    response_texts = ask_model(row["title"], row["abstract"], row["methods"], row["results"])
    all_answers = parse_answers(response_texts[0])

    remaining_df.at[index, f"sample_1_raw_response"] = clean_text(response_texts[0])

    for q_idx in range(1, 4):
        remaining_df.at[index, f"sample_1_q{q_idx}_answer"] = all_answers[q_idx - 1]["answer"]
        remaining_df.at[index, f"sample_1_q{q_idx}_reasoning"] = clean_text(all_answers[q_idx - 1]["reasoning"])

    full_row = remaining_df.loc[[index]]

    full_row.to_csv(
        LLM_LABELED_DATA_PATH,
        mode='a',
        header=not os.path.exists(LLM_LABELED_DATA_PATH),
        index=False,
        sep=";",
        quoting=csv.QUOTE_MINIMAL
    )

print(f"\nBatch processing complete. Processed {min(NUM_ARTICLES_TO_PROCESS, len(remaining_df))} articles.")
print(f"Results saved to {LLM_LABELED_DATA_PATH}")
```
