## Supplementary material for "Easy-to-use whole-genome sequencing workflows and standardized practices to uncover hidden genetic variation in *Synechocystis* PCC 6803 wild-type and knock-out strains": S8 File: INDEL_generation.py

```
#!/usr/bin/env python3

from pathlib import Path
import random
import argparse
from collections import Counter, defaultdict

# ---------- Defaults ----------
INPUT_FASTA = "Hess_Genome.fasta"
OUTPUT_FASTA = "Hess_Genome_INDELS_1.fasta"
MUTATION_TSV = "Hess_Genome_INDELS_All_1.tsv"
VCF_OUT = "Hess_Genome_INDELS_All_1.vcf"

NUM_RANDOM_INDELS = 40
NUM_REPEAT_INDELS = 10
MIN_REPEAT = 6

MIN_SIZE = -10
MAX_SIZE = 10

BUFFER_SIZE = 3500
SEED = 4
ALLOW_N = False
MAX_TRIES_FACTOR = 5
# ---------------------------------------------------------------


def read_fasta(path):
    contigs, h, buf = {}, None, []
    with open(path) as f:
        for line in f:
            s = line.rstrip()
            if not s:
                continue
            if s.startswith(">"):
                if h:
                    contigs[h] = "".join(buf).upper()
                h, buf = s[1:].strip(), []
            else:
                buf.append(s)
        if h:
            contigs[h] = "".join(buf).upper()
    print(f"Loaded {len(contigs)} contigs, total {sum(len(s) for s in contigs.values()):,} bp")
    return contigs


def write_fasta(path, contigs, width=60):
    with open(path, "w") as f:
        for h, s in contigs.items():
            f.write(f">{h}\n")
            for i in range(0, len(s), width):
                f.write(s[i:i+width] + "\n")
    print(f"Wrote FASTA: {path}")


def write_tsv(path, muts):
    with open(path, "w") as f:
        f.write("contig\tpos_1based\tsize\ttype\tref\talt\tindel_type\n")
        for m in muts:
            f.write(
                f"{m['contig']}\t{m['pos_1based']}\t{m['size']}\t"
                f"{m['type']}\t{m['ref']}\t{m['alt']}\t{m['indel_type']}\n"
            )
    print(f"Wrote TSV: {path} ({len(muts)} records)")


def write_vcf(path, muts, contigs_ref, sample_name="SAMPLE"):
    """
    Write planned mutations to a VCF using ORIGINAL reference coordinates.

    Insertions:
      POS = p+1 (same as muts pos_1based)
      REF = anchor base
      ALT = anchor base + inserted bases

    Deletions:
      Use left anchor base (preceding base):
      POS = anchor_position (1-based)
      REF = anchor_base + deleted_bases
      ALT = anchor_base
    """
    contig_lines = [f"##contig=<ID={h},length={len(seq)}>" for h, seq in contigs_ref.items()]

    with open(path, "w") as f:
        f.write("##fileformat=VCFv4.2\n")
        f.write("##source=INDEL_mutator\n")
        for line in contig_lines:
            f.write(line + "\n")
        f.write('##INFO=<ID=SVTYPE,Number=1,Type=String,Description="Variant type">\n')
        f.write('##INFO=<ID=SVLEN,Number=1,Type=Integer,Description="Length of variant (ALT-REF)">\n')
        f.write('##INFO=<ID=INDELTYPE,Number=1,Type=String,Description="random|repeat">\n')
        f.write('##FORMAT=<ID=GT,Number=1,Type=String,Description="Genotype">\n')
        f.write("#CHROM\tPOS\tID\tREF\tALT\tQUAL\tFILTER\tINFO\tFORMAT\t" + sample_name + "\n")

        for i, m in enumerate(sorted(muts, key=lambda x: (x["contig"], x["pos_1based"]))):
            chrom = m["contig"]
            refseq = contigs_ref[chrom]
            p0 = m["pos_1based"] - 1  # 0-based in original reference

            if m["type"] == "INS":
                pos = m["pos_1based"]
                ref = m["ref"]
                alt = m["alt"]
                svtype = "INS"
                svlen = len(alt) - len(ref)
            else:  # DEL
                d = abs(m["size"])
                if p0 == 0:
                    raise ValueError(
                        f"Cannot represent deletion at start of contig in VCF without a left anchor: {chrom}:{m['pos_1based']}"
                    )
                anchor0 = p0 - 1
                anchor_base = refseq[anchor0]
                deleted = refseq[p0:p0 + d]

                pos = anchor0 + 1
                ref = anchor_base + deleted
                alt = anchor_base
                svtype = "DEL"
                svlen = -d

            info = f"SVTYPE={svtype};SVLEN={svlen};INDELTYPE={m['indel_type']}"
            vid = f"indel{i+1}"
            f.write(f"{chrom}\t{pos}\t{vid}\t{ref}\t{alt}\t.\tPASS\t{info}\tGT\t1/1\n")

    print(f"Wrote VCF: {path} ({len(muts)} records)")


def make_even_size_list(n, lo, hi):
    sizes = [i for i in range(lo, hi + 1) if i != 0]
    out = []
    while len(out) < n:
        random.shuffle(sizes)
        out.extend(sizes)
    return out[:n]


def find_homopolymers(seq, min_len):
    out, i, L = [], 0, len(seq)
    while i < L:
        b, j = seq[i], i + 1
        while j < L and seq[j] == b:
            j += 1
        if b in "ACGT" and j - i >= min_len:
            out.append((i, b))
        i = j
    random.shuffle(out)
    return out


def reserve_buffer(bufs, h, s, e, bs):
    r = range(max(0, s - bs), e + bs + 1)
    if any(p in bufs[h] for p in r):
        return False
    bufs[h].update(r)
    return True


def region_is_acgt(seq, s, e):
    return all(b in "ACGT" for b in seq[s:e+1])


def plan_repeat_indels(contigs, n, min_rep, bufs, bs):
    cands = [(h, p, b)
             for h, s in contigs.items()
             for p, b in find_homopolymers(s, min_rep)]
    random.shuffle(cands)

    ops = [1] * (n // 2) + [-1] * (n - n // 2)
    random.shuffle(ops)

    base_cap = max(1, n // 4)
    base_cnt = Counter()
    out = []

    i = 0
    for h, p, b in cands:
        if i >= n:
            break
        if base_cnt[b] >= base_cap:
            continue
        if not reserve_buffer(bufs, h, p, p, bs):
            continue

        size = ops[i]
        run_end = p
        while run_end + 1 < len(contigs[h]) and contigs[h][run_end + 1] == b:
            run_end += 1

        out.append(dict(
            contig=h,
            ref_pos=run_end,
            size=size,
            type="INS" if size > 0 else "DEL",
            indel_type="repeat"
        ))

        base_cnt[b] += 1
        i += 1

        if len(out) == n:
            break

    print(f"Planned repeat indels: {len(out)}/{n}")
    return out


def plan_random_indels(contigs, n, sizes, allow_n, bufs, bs):
    hs = list(contigs)
    lens = {h: len(contigs[h]) for h in hs}
    wts = [lens[h] for h in hs]

    out, tries = [], 0
    max_tries = MAX_TRIES_FACTOR * n * len(hs)

    for sz in sizes:
        if len(out) == n or tries > max_tries:
            break
        tries += 1
        for _ in range(100):
            h = random.choices(hs, wts)[0]
            L = lens[h]
            p = random.randint(0, L - 1)

            s, e = (p, p) if sz > 0 else (p, p + abs(sz) - 1)
            if e >= L:
                continue
            if not allow_n and not region_is_acgt(contigs[h], s, e):
                continue
            if not reserve_buffer(bufs, h, s, e, bs):
                continue

            out.append(dict(contig=h, ref_pos=p, size=sz,
                            type="INS" if sz > 0 else "DEL",
                            indel_type="random"))
            break

    print(f"Planned random indels: {len(out)}/{n}")
    return out


def apply_mutations(contigs, mutations):
    by_contig = defaultdict(list)
    for m in mutations:
        by_contig[m["contig"]].append(m)

    for h, muts in by_contig.items():
        orig = contigs[h]
        seq = list(orig)

        for m in sorted(muts, key=lambda x: x["ref_pos"], reverse=True):
            p = m["ref_pos"]

            if m["size"] < 0:
                d = abs(m["size"])
                m["ref"] = orig[p:p+d]
                m["alt"] = ""
                del seq[p:p+d]
            else:
                a = orig[p]
                ins = a * m["size"] if m["indel_type"] == "repeat" \
                    else "".join(random.choice("ACGT") for _ in range(m["size"]))
                m["ref"] = a
                m["alt"] = a + ins
                seq[p+1:p+1] = list(ins)

            m["pos_1based"] = p + 1

        contigs[h] = "".join(seq)

    return contigs


def main(args):
    random.seed(args.seed)
    if not Path(args.input).exists():
        raise FileNotFoundError(args.input)

    contigs = read_fasta(args.input)
    contigs_ref = dict(contigs)  # keep ORIGINAL reference for VCF anchoring
    bufs = {h: set() for h in contigs}

    muts = (
        plan_repeat_indels(contigs, args.repeat, args.min_repeat, bufs, args.buffer_size)
        + plan_random_indels(
            contigs, args.random,
            make_even_size_list(args.random, args.min_size, args.max_size),
            args.allow_n, bufs, args.buffer_size)
    )

    if not muts:
        print("No mutations planned")
        return

    write_fasta(args.output, apply_mutations(contigs, muts))
    write_tsv(args.tsv, muts)
    write_vcf(args.vcf, muts, contigs_ref)
    print(f"Done. Total indels: {len(muts)}")


if __name__ == "__main__":
    p = argparse.ArgumentParser()
    p.add_argument("--in", dest="input", default=INPUT_FASTA)
    p.add_argument("--out", dest="output", default=OUTPUT_FASTA)
    p.add_argument("--tsv", dest="tsv", default=MUTATION_TSV)
    p.add_argument("--vcf", dest="vcf", default=VCF_OUT)

    p.add_argument("--random", type=int, default=NUM_RANDOM_INDELS)
    p.add_argument("--repeat", type=int, default=NUM_REPEAT_INDELS)
    p.add_argument("--min-repeat", type=int, default=MIN_REPEAT)
    p.add_argument("--min-size", type=int, default=MIN_SIZE)
    p.add_argument("--max-size", type=int, default=MAX_SIZE)
    p.add_argument("--buffer-size", type=int, default=BUFFER_SIZE)
    p.add_argument("--seed", type=int, default=SEED)
    p.add_argument("--allow-n", action="store_true", default=ALLOW_N)

    main(p.parse_args())
```
