## Supplementary material for "Easy-to-use whole-genome sequencing workflows and standardized practices to uncover hidden genetic variation in *Synechocystis* PCC 6803 wild-type and knock-out strains": S8 File: RelDup_Achor_fixed.py

```
#!/usr/bin/env python3

from pathlib import Path
import random
import argparse
from datetime import datetime


# ---------------- FASTA I/O ----------------

def read_fasta(path):
    contigs = {}
    header = None
    seq_lines = []
    with open(path) as f:
        for line in f:
            s = line.rstrip()
            if not s:
                continue
            if s.startswith(">"):
                if header:
                    contigs[header] = "".join(seq_lines).upper()
                header = s[1:].strip()
                seq_lines = []
            else:
                seq_lines.append(s)
        if header:
            contigs[header] = "".join(seq_lines).upper()
    return contigs


def write_fasta(path, contigs, wrap=80):
    with open(path, "w") as f:
        for h, s in contigs.items():
            f.write(f">{h}\n")
            if wrap and wrap > 0:
                for i in range(0, len(s), wrap):
                    f.write(s[i:i+wrap] + "\n")
            else:
                f.write(s + "\n")


# ---------------- OLD BUFFER LOGIC (unchanged) ----------------

def reserve_buffer(buffer_sets, contig, start, end, buffer_size):
    reserved = buffer_sets[contig]
    rs = max(0, start - buffer_size)
    re = end + buffer_size
    for p in range(rs, re + 1):
        if p in reserved:
            return False
    for p in range(rs, re + 1):
        reserved.add(p)
    return True


# ---------------- PLANNING (unchanged behavior/order) ----------------

def plan_reldup_svs_like_old(contigs, sizes, num_copies, buffer_sets, buffer_size, seed, allow_intercontig=True):
    random.seed(seed)

    svs = []
    contig_names = list(contigs.keys())

    for sv_type in ["Relocation", "Duplication"]:
        for size in sizes:
            for _ in range(num_copies):
                tries = 0
                max_tries = 1000
                while tries < max_tries:
                    tries += 1

                    contig_lens_dict = {h: len(contigs[h]) for h in contig_names}
                    contig_lens_list = [contig_lens_dict[h] for h in contig_names]

                    src_contig = random.choices(contig_names, weights=contig_lens_list, k=1)[0]
                    L = contig_lens_dict[src_contig]
                    if size >= L:
                        continue

                    src_start = random.randint(0, L - size)
                    src_end = src_start + size - 1

                    if not reserve_buffer(buffer_sets, src_contig, src_start, src_end, buffer_size):
                        continue

                    tgt_contig = random.choices(contig_names, weights=contig_lens_list, k=1)[0]
                    if (not allow_intercontig) and (tgt_contig != src_contig):
                        continue

                    tgt_len = contig_lens_dict[tgt_contig]
                    tgt_pos = random.randint(0, tgt_len)

                    if not reserve_buffer(buffer_sets, tgt_contig, tgt_pos, tgt_pos, buffer_size):
                        continue

                    svs.append({
                        "sv_id": f"SV{len(svs)+1:05d}",
                        "type": sv_type,
                        "src_contig": src_contig,
                        "src_start": src_start,
                        "src_end": src_end,
                        "dst_contig": tgt_contig,
                        "dst_pos": tgt_pos,
                        "size": size,
                    })
                    break

                if tries >= max_tries:
                    raise RuntimeError(f"Could not place {sv_type} size {size} after {max_tries} tries")

    return svs


# ---------------- Signature capture (around breakpoint) ----------------

def get_signature(seq, breakpoint_pos0, up_len, down_len):
    """
    Return signature string spanning [breakpoint-up_len : breakpoint+down_len)
    clipped to contig bounds. Also returns actual upstream length used (may be smaller near start).
    """
    bp = max(0, min(len(seq), breakpoint_pos0))
    left = max(0, bp - up_len)
    right = min(len(seq), bp + down_len)
    sig = seq[left:right]
    actual_up = bp - left
    return sig, actual_up


# ---------------- APPLY (mutation logic unchanged) + capture signatures ----------------

def apply_reldup_svs_like_old_and_capture(contigs, svs, up_len=100, down_len=30):
    """
    Applies SVs EXACTLY like old script.
    Additionally captures a breakpoint signature (upstream+downstream) at:
      - relocation deletion breakpoint: src_start (before deletion)
      - relocation insertion breakpoint: dst_pos (after deletion, before insertion)
      - duplication insertion breakpoint: dst_pos (before insertion)
    """
    for m in svs:
        src_c = m["src_contig"]
        src_s = m["src_start"]
        src_e = m["src_end"]
        dst_c = m["dst_contig"]
        dst_p = m["dst_pos"]

        # payload taken from CURRENT contig at apply-time (old behavior)
        payload = contigs[src_c][src_s:src_e+1]

        if m["type"] == "Relocation":
            # deletion breakpoint signature (before deletion)
            sig, actual_up = get_signature(contigs[src_c], src_s, up_len, down_len)
            m["del_sig_contig"] = src_c
            m["del_sig"] = sig
            m["del_sig_up"] = actual_up  # usually up_len, smaller near contig start

            # delete (unchanged)
            contigs[src_c] = contigs[src_c][:src_s] + contigs[src_c][src_e+1:]

            # insertion breakpoint signature (after deletion, before insertion)
            sig, actual_up = get_signature(contigs[dst_c], dst_p, up_len, down_len)
            m["ins_sig_contig"] = dst_c
            m["ins_sig"] = sig
            m["ins_sig_up"] = actual_up

            # insert (unchanged)
            contigs[dst_c] = contigs[dst_c][:dst_p] + payload + contigs[dst_c][dst_p:]

        elif m["type"] == "Duplication":
            # insertion breakpoint signature (before insertion)
            sig, actual_up = get_signature(contigs[dst_c], dst_p, up_len, down_len)
            m["ins_sig_contig"] = dst_c
            m["ins_sig"] = sig
            m["ins_sig_up"] = actual_up

            # insert (unchanged)
            contigs[dst_c] = contigs[dst_c][:dst_p] + payload + contigs[dst_c][dst_p:]
        else:
            raise ValueError(m["type"])

    return contigs


# ---------------- Map signatures back onto ORIGINAL reference ----------------

def find_all(haystack, needle):
    hits = []
    i = haystack.find(needle)
    while i != -1:
        hits.append(i)
        i = haystack.find(needle, i + 1)
    return hits


def locate_signature_on_original(original_contigs, contig, sig, hint_bp0=None):
    """
    Return start index (0-based) of signature in original contig.
    If multiple hits: choose closest to (hint_bp0 - up_used) i.e. closest expected neighborhood.
    """
    seq = original_contigs[contig]
    hits = find_all(seq, sig)
    if not hits:
        raise ValueError(f"Signature not found on ORIGINAL: {contig} (len(sig)={len(sig)})")
    if len(hits) == 1:
        return hits[0]
    if hint_bp0 is None:
        raise ValueError(f"Signature not unique on ORIGINAL: {contig} (hits={len(hits)})")
    return min(hits, key=lambda h: abs(h - hint_bp0))


def map_breakpoints_to_original(original_contigs, svs):
    """
    Computes ORIGINAL-coordinate breakpoints from stored signatures.
    Breakpoint (0-based) = signature_match_start + stored_actual_up
    Hints:
      - deletion breakpoint hint uses planned src_start
      - insertion breakpoint hint uses planned dst_pos
    """
    for m in svs:
        if m["type"] == "Relocation":
            # deletion breakpoint on original
            c = m["del_sig_contig"]
            sig = m["del_sig"]
            up_used = m["del_sig_up"]
            hint = max(0, m["src_start"] - up_used)
            start0 = locate_signature_on_original(original_contigs, c, sig, hint_bp0=hint)
            m["orig_del_bp0"] = start0 + up_used

            # insertion breakpoint on original
            c = m["ins_sig_contig"]
            sig = m["ins_sig"]
            up_used = m["ins_sig_up"]
            hint = max(0, m["dst_pos"] - up_used)
            start0 = locate_signature_on_original(original_contigs, c, sig, hint_bp0=hint)
            m["orig_ins_bp0"] = start0 + up_used

        elif m["type"] == "Duplication":
            c = m["ins_sig_contig"]
            sig = m["ins_sig"]
            up_used = m["ins_sig_up"]
            hint = max(0, m["dst_pos"] - up_used)
            start0 = locate_signature_on_original(original_contigs, c, sig, hint_bp0=hint)
            m["orig_ins_bp0"] = start0 + up_used
        else:
            raise ValueError(m["type"])


# ---------------- TSV / VCF (ORIGINAL coords derived from signature mapping) ----------------

def write_tsv(path, svs):
    header = ["sv_id", "event", "contig", "pos_1based", "size", "description"]
    with open(path, "w") as f:
        f.write("\t".join(header) + "\n")
        for m in svs:
            if m["type"] == "Relocation":
                # Deletion breakpoint (source)
                f.write("\t".join(map(str, [
                    m["sv_id"],
                    "Relocation_DEL",
                    m["src_contig"],
                    m["orig_del_bp0"] + 1,
                    m["size"],
                    f"Deleted {m['size']}bp from {m['src_contig']} at {m['orig_del_bp0']+1} (ORIGINAL via signature)"
                ])) + "\n")

                # Insertion breakpoint (destination)
                f.write("\t".join(map(str, [
                    m["sv_id"],
                    "Relocation_INS",
                    m["dst_contig"],
                    m["orig_ins_bp0"] + 1,
                    m["size"],
                    f"Inserted {m['size']}bp into {m['dst_contig']} at {m['orig_ins_bp0']+1} (ORIGINAL via signature)"
                ])) + "\n")

            elif m["type"] == "Duplication":
                f.write("\t".join(map(str, [
                    m["sv_id"],
                    "Duplication_INS",
                    m["dst_contig"],
                    m["orig_ins_bp0"] + 1,
                    m["size"],
                    f"Inserted duplicated {m['size']}bp into {m['dst_contig']} at {m['orig_ins_bp0']+1} (ORIGINAL via signature)"
                ])) + "\n")
            else:
                raise ValueError(m["type"])

def _vcf_escape(s):
    return str(s).replace("\\", "\\\\").replace("\t", " ").replace("\n", " ")


def write_vcf(path, original_contigs, svs):
    with open(path, "w") as f:
        dt = datetime.now().strftime("%Y%m%d")
        f.write("##fileformat=VCFv4.2\n")
        f.write(f"##fileDate={dt}\n")
        f.write("##source=SVMutator_signature_remap_to_ORIGINAL\n")
        f.write('##ALT=<ID=DUP,Description="Duplication">\n')
        f.write('##ALT=<ID=BND,Description="Breakend">\n')
        f.write('##INFO=<ID=SVTYPE,Number=1,Type=String,Description="Type of structural variant">\n')
        f.write('##INFO=<ID=SVLEN,Number=1,Type=Integer,Description="Length of the SV">\n')
        f.write('##INFO=<ID=MATEID,Number=1,Type=String,Description="ID of mate breakend">\n')
        for c, s in original_contigs.items():
            f.write(f"##contig=<ID={_vcf_escape(c)},length={len(s)}>\n")
        f.write("#CHROM\tPOS\tID\tREF\tALT\tQUAL\tFILTER\tINFO\n")

        for m in svs:
            size = m["size"]

            if m["type"] == "Duplication":
                chrom = m["dst_contig"]
                pos = m["orig_ins_bp0"] + 1
                f.write(
                    f"{chrom}\t{pos}\t{m['sv_id']}\tN\t<DUP>\t.\tPASS\t"
                    f"SVTYPE=DUP;SVLEN={size}\n"
                )

            elif m["type"] == "Relocation":
                chrom1 = m["src_contig"]
                pos1 = m["orig_del_bp0"] + 1
                chrom2 = m["dst_contig"]
                pos2 = m["orig_ins_bp0"] + 1

                id1 = m["sv_id"] + "_A"
                id2 = m["sv_id"] + "_B"
                alt1 = f"N]{chrom2}:{pos2}]"
                alt2 = f"N]{chrom1}:{pos1}]"

                f.write(
                    f"{chrom1}\t{pos1}\t{id1}\tN\t{alt1}\t.\tPASS\t"
                    f"SVTYPE=BND;SVLEN={size};MATEID={id2}\n"
                )
                f.write(
                    f"{chrom2}\t{pos2}\t{id2}\tN\t{alt2}\t.\tPASS\t"
                    f"SVTYPE=BND;SVLEN={size};MATEID={id1}\n"
                )
            else:
                raise ValueError(m["type"])


# ---------------- MAIN ----------------

def main():
    ap = argparse.ArgumentParser(description="SV mutator (old behavior) + signature remap to ORIGINAL coords.")
    ap.add_argument("--input", default="Hess_Genome.fasta")
    ap.add_argument("--output-fasta", default="Hess_Genome_RelDup_SV_fixed_5.fasta")
    ap.add_argument("--output-tsv", default="Hess_Genome_RelDup_SV_fixed_5.tsv")
    ap.add_argument("--output-vcf", default="Hess_Genome_RelDup_SV_fixed_5.vcf")

    ap.add_argument("--seed", type=int, default=32)
    ap.add_argument("--buffer-size", type=int, default=3500)
    ap.add_argument("--allow-intercontig", action="store_true", default=True)

    ap.add_argument("--sv-sizes", type=int, nargs="+", default=[100, 200, 500, 1000, 2000])
    ap.add_argument("--num-copies", type=int, default=3)

    ap.add_argument("--up-len", type=int, default=100)
    ap.add_argument("--down-len", type=int, default=30)
    ap.add_argument("--fasta-wrap", type=int, default=80)

    args = ap.parse_args()

    if not Path(args.input).exists():
        raise FileNotFoundError(args.input)

    original = read_fasta(args.input)

    buffer_sets = {h: set() for h in original}
    svs = plan_reldup_svs_like_old(
        contigs=original,
        sizes=args.sv_sizes,
        num_copies=args.num_copies,
        buffer_sets=buffer_sets,
        buffer_size=args.buffer_size,
        seed=args.seed,
        allow_intercontig=args.allow_intercontig,
    )

    mutated = {h: s for h, s in original.items()}
    mutated = apply_reldup_svs_like_old_and_capture(mutated, svs, up_len=args.up_len, down_len=args.down_len)

    map_breakpoints_to_original(original, svs)

    write_fasta(args.output_fasta, mutated, wrap=args.fasta_wrap)
    write_tsv(args.output_tsv, svs)
    write_vcf(args.output_vcf, original, svs)

if __name__ == "__main__":
    main()
```
