## Supplementary material for "Easy-to-use whole-genome sequencing workflows and standardized practices to uncover hidden genetic variation in *Synechocystis* PCC 6803 wild-type and knock-out strains": S8 File: SNP_generator_2.py

```
#!/usr/bin/env python3

from pathlib import Path
import random
import argparse

# ---------- Defaults ----------
INPUT_FASTA = "Hess_Genome.fasta"
OUTPUT_FASTA = "Hess_Genome_SNPs_5.fasta"
MUTATION_TSV = "Hess_Genome_SNPs_5.tsv"
VCF_OUT = "Hess_Genome_SNPs_5.vcf"

NUM_RANDOM_SNPS = 40
NUM_REPEAT_SNPS = 10
MIN_REPEAT = 6

BUFFER_SIZE = 3500
SEED = 4
ALLOW_N = False
MAX_TRIES_FACTOR = 5
# ---------------------------------------------------------------

def read_fasta(path):
    contigs = {}
    header = None
    seq_lines = []
    with open(path) as f:
        for line in f:
            s = line.rstrip()
            if not s:
                continue
            if s.startswith(">"):
                if header:
                    contigs[header] = "".join(seq_lines).upper()
                header = s[1:].strip()
                seq_lines = []
            else:
                seq_lines.append(s)
        if header:
            contigs[header] = "".join(seq_lines).upper()
    print(f"Loaded {len(contigs)} contigs, total {sum(len(s) for s in contigs.values()):,} bp")
    return contigs

def write_fasta(path, contigs):
    with open(path, "w") as f:
        for h, s in contigs.items():
            f.write(f">{h}\n")
            f.write(s + "\n")
    print(f"Wrote FASTA: {path}")

def write_tsv(path, muts):
    with open(path, "w") as f:
        f.write("contig\tposition_1based\tref\talt\tmutation_type\textra_info\n")
        for m in muts:
            f.write(f"{m['contig']}\t{m['position_1based']}\t{m['ref']}\t{m['alt']}\t{m['mutation_type']}\t{m['extra_info']}\n")
    print(f"Wrote TSV: {path} ({len(muts)} records)")

def write_vcf(path, muts, contigs_ref, sample_name="SAMPLE"):
    """
    Write SNPs to VCF (VCFv4.2) in ORIGINAL reference coordinates.
    Uses planned ref/alt in muts; does not change any mutation logic.
    """
    contig_lines = [f"##contig=<ID={h},length={len(seq)}>" for h, seq in contigs_ref.items()]

    with open(path, "w") as f:
        f.write("##fileformat=VCFv4.2\n")
        f.write("##source=SNP_mutator\n")
        for line in contig_lines:
            f.write(line + "\n")
        f.write('##INFO=<ID=MUTTYPE,Number=1,Type=String,Description="repeat|random">\n')
        f.write('##INFO=<ID=EXTRA,Number=1,Type=String,Description="extend_repeat|shorten_repeat|-">\n')
        f.write('##FORMAT=<ID=GT,Number=1,Type=String,Description="Genotype">\n')
        f.write("#CHROM\tPOS\tID\tREF\tALT\tQUAL\tFILTER\tINFO\tFORMAT\t" + sample_name + "\n")

        # Ensure position_1based exists (it is set in apply_snps; but compute if missing)
        for i, m in enumerate(sorted(muts, key=lambda x: (x["contig"], x.get("position_1based", x["pos"] + 1)))):
            chrom = m["contig"]
            pos = m.get("position_1based", m["pos"] + 1)
            ref = m["ref"]
            alt = m["alt"]

            # Minimal sanity: VCF REF/ALT for SNP should be 1 base (A/C/G/T/N)
            # Keep as-is to avoid logic changes.
            vid = f"snp{i+1}"
            info = f"MUTTYPE={m['mutation_type']};EXTRA={m['extra_info']}"
            f.write(f"{chrom}\t{pos}\t{vid}\t{ref}\t{alt}\t.\tPASS\t{info}\tGT\t1/1\n")

    print(f"Wrote VCF: {path} ({len(muts)} records)")

def reserve_buffer(buffer_sets, contig, pos, buffer_size):
    """Return True and reserve if free; otherwise False."""
    reserved = buffer_sets[contig]
    rs = max(0, pos - buffer_size)
    re = pos + buffer_size
    for p in range(rs, re+1):
        if p in reserved:
            return False
    for p in range(rs, re+1):
        reserved.add(p)
    return True

def find_homopolymers(seq, min_len):
    """Return list of (start, base, length) for runs >= min_len."""
    sites = []
    i = 0
    L = len(seq)
    while i < L:
        b = seq[i]
        j = i + 1
        while j < L and seq[j] == b:
            j += 1
        run = j - i
        if b in "ACGT" and run >= min_len:
            sites.append((i, b, run))
        i = j
    random.shuffle(sites)
    return sites

def plan_repeat_snps(contigs, num_repeat, min_repeat, buffer_sets):
    """Plan SNPs at homopolymers using INDEL-style placement."""
    candidates = []
    for h, seq in contigs.items():
        for start, base, run in find_homopolymers(seq, min_repeat):
            # allow positions inside or immediately after the run
            for offset in range(run + 1):
                pos = start + offset
                if pos >= len(seq):
                    continue
                candidates.append((h, pos, base, seq[pos]))

    random.shuffle(candidates)
    n = min(len(candidates), num_repeat)
    half = n // 2
    planned = []
    i = 0

    for h, pos, base, ref in candidates:
        if i >= n:
            break
        if not reserve_buffer(buffer_sets, h, pos, BUFFER_SIZE):
            continue
        # extend repeat
        if i < half:
            if ref == base:
                continue
            alt = base
            extra = "extend_repeat"
        # shorten repeat
        else:
            choices = [b for b in "ACGT" if b != base]
            if not choices or ref in choices:
                continue
            alt = random.choice(choices)
            extra = "shorten_repeat"
        planned.append({
            "contig": h,
            "pos": pos,
            "ref": ref,
            "alt": alt,
            "mutation_type": "repeat",
            "extra_info": extra
        })
        i += 1

    print(f"Planned repeat SNPs: {len(planned)}/{num_repeat}")
    return planned

def plan_random_snps(contigs, num_snps, avoid_n, buffer_sets):
    """Plan random SNPs across contigs using INDEL-style placement."""
    planned = []
    contig_names = list(contigs.keys())
    contig_lens = {h: len(contigs[h]) for h in contig_names}
    weights = [contig_lens[h] for h in contig_names]

    max_global_tries = MAX_TRIES_FACTOR * num_snps * len(contig_names)
    tries = 0

    while len(planned) < num_snps and tries < max_global_tries:
        tries += 1
        h = random.choices(contig_names, weights=weights, k=1)[0]
        L = contig_lens[h]
        if L == 0:
            continue
        for _ in range(100):
            pos = random.randint(0, L-1)
            base = contigs[h][pos]
            if avoid_n and base not in "ACGT":
                continue
            if not reserve_buffer(buffer_sets, h, pos, BUFFER_SIZE):
                continue
            alt_choices = [b for b in "ACGT" if b != base]
            if not alt_choices:
                continue
            alt = random.choice(alt_choices)
            planned.append({
                "contig": h,
                "pos": pos,
                "ref": base,
                "alt": alt,
                "mutation_type": "random",
                "extra_info": "-"
            })
            break

    if len(planned) < num_snps:
        print(f"Warning: Only placed {len(planned)}/{num_snps} random SNPs after {tries} tries")
    else:
        print(f"Planned {len(planned)} random SNPs successfully")
    return planned

def apply_snps(contigs, snps):
    """Apply SNPs to contigs."""
    for m in snps:
        h = m["contig"]
        pos = m["pos"]
        seq = list(contigs[h])
        seq[pos] = m["alt"]
        contigs[h] = "".join(seq)
        m["position_1based"] = pos + 1
    return contigs

def main(args):
    random.seed(args.seed)
    if not Path(args.input).exists():
        raise FileNotFoundError(args.input)

    contigs = read_fasta(args.input)
    contigs_ref = dict(contigs)  # keep original reference for VCF header/lengths
    buffer_sets = {h: set() for h in contigs.keys()}

    repeat_snps = plan_repeat_snps(contigs, args.repeat, args.min_repeat, buffer_sets)
    random_snps = plan_random_snps(contigs, args.random, not args.allow_n, buffer_sets)

    all_snps = repeat_snps + random_snps
    if not all_snps:
        print("No SNPs planned.")
        return

    mutated = apply_snps(contigs, all_snps)
    write_fasta(args.output, mutated)
    write_tsv(args.tsv, all_snps)
    write_vcf(args.vcf, all_snps, contigs_ref)

    print(f"Done. Total SNPs: {len(all_snps)} "
          f"(repeat={len(repeat_snps)}, random={len(random_snps)})")

if __name__ == "__main__":
    p = argparse.ArgumentParser()
    p.add_argument("--in", dest="input", default=INPUT_FASTA)
    p.add_argument("--out", dest="output", default=OUTPUT_FASTA)
    p.add_argument("--tsv", dest="tsv", default=MUTATION_TSV)
    p.add_argument("--vcf", dest="vcf", default=VCF_OUT)

    p.add_argument("--random", type=int, default=NUM_RANDOM_SNPS)
    p.add_argument("--repeat", type=int, default=NUM_REPEAT_SNPS)
    p.add_argument("--min-repeat", type=int, default=MIN_REPEAT)
    p.add_argument("--buffer-size", type=int, default=BUFFER_SIZE)
    p.add_argument("--seed", type=int, default=SEED)
    p.add_argument("--allow-n", action="store_true", default=False)
    args = p.parse_args()
    globals()["BUFFER_SIZE"] = args.buffer_size

    main(args)
```
