## Supplementary material for "Easy-to-use whole-genome sequencing workflows and standardized practices to uncover hidden genetic variation in *Synechocystis* PCC 6803 wild-type and knock-out strains": S8 File: SV_Insertion_Deletion.py

```
#!/usr/bin/env python3

from pathlib import Path
import random
import argparse
from collections import defaultdict
from datetime import datetime

# ---------- Defaults ----------
INPUT_FASTA = "Hess_Genome.fasta"
OUTPUT_FASTA = "Hess_Genome_InDel_All_1.fasta"
MUTATION_TSV = "Hess_Genome_InDel_All_1.tsv"
OUTPUT_VCF = "Hess_Genome_InDel_All_1.vcf"

BUFFER_SIZE = 3500
SEED =2

# SV sizes and counts
SV_SIZES = [100, 200, 500, 1000, 2000]
NUM_COPIES = 1  # 10 of each size for both insertions and deletions

# ---------------------------------------------------------------

def read_fasta(path):
    """Read FASTA file into dictionary"""
    contigs = {}
    header = None
    seq_lines = []
    with open(path) as f:
        for line in f:
            s = line.rstrip()
            if not s:
                continue
            if s.startswith(">"):
                if header:
                    contigs[header] = "".join(seq_lines).upper()
                header = s[1:].strip()
                seq_lines = []
            else:
                seq_lines.append(s)
        if header:
            contigs[header] = "".join(seq_lines).upper()
    print(f"Loaded {len(contigs)} contigs, total {sum(len(s) for s in contigs.values()):,} bp")
    return contigs

def write_fasta(path, contigs):
    """Write contigs to FASTA file"""
    with open(path, "w") as f:
        for h, s in contigs.items():
            f.write(f">{h}\n")
            f.write(s + "\n")
    print(f"Wrote FASTA: {path}")

def write_tsv(path, truth_records):
    """Write truth table with reference genome coordinates"""
    with open(path, "w") as f:
        f.write("contig\tstart_1based\tend_1based\ttype\tsize\tdescription\n")
        for rec in truth_records:
            f.write(f"{rec['contig']}\t{rec['start_1based']}\t{rec['end_1based']}\t"
                   f"{rec['type']}\t{rec['size']}\t{rec['description']}\n")
    print(f"Wrote TSV: {path} ({len(truth_records)} records)")

def generate_random_sequence(length):
    """Generate a random DNA sequence of given length"""
    bases = ['A', 'C', 'G', 'T']
    return ''.join(random.choice(bases) for _ in range(length))

def check_buffer_overlap(reserved_regions, contig, start, end, buffer_size):
    """Check if region overlaps with any reserved region (including buffer)"""
    if contig not in reserved_regions:
        return False
    
    check_start = max(0, start - buffer_size)
    check_end = end + buffer_size
    
    for res_start, res_end in reserved_regions[contig]:
        # Check for any overlap
        if not (check_end < res_start or check_start > res_end):
            return True
    return False

def reserve_region(reserved_regions, contig, start, end, buffer_size):
    """Reserve a region with buffer"""
    if contig not in reserved_regions:
        reserved_regions[contig] = []
    
    reserve_start = max(0, start - buffer_size)
    reserve_end = end + buffer_size
    reserved_regions[contig].append((reserve_start, reserve_end))

def plan_all_svs(original_contigs, sizes, num_copies, buffer_size, seed):
    """
    Plan ALL structural variants before applying any mutations.
    All coordinates are relative to the ORIGINAL reference genome.
    """
    random.seed(seed)
    
    # Make a copy to check lengths without mutating
    contig_names = list(original_contigs.keys())
    contig_lengths = {h: len(original_contigs[h]) for h in contig_names}
    
    # Track reserved regions to avoid overlaps
    reserved_regions = defaultdict(list)
    
    sv_plans = []
    
    # Plan deletions
    print("Planning deletions...")
    for size in sizes:
        placed = 0
        attempts = 0
        max_attempts = 10000
        
        while placed < num_copies and attempts < max_attempts:
            attempts += 1
            
            # Pick contig (weighted by length)
            contig = random.choices(contig_names, 
                                   weights=[contig_lengths[h] for h in contig_names])[0]
            contig_len = contig_lengths[contig]
            
            if size >= contig_len:
                continue
                
            # Pick deletion position
            del_start = random.randint(0, contig_len - size)
            del_end = del_start + size - 1
            
            # Check if region is available
            if check_buffer_overlap(reserved_regions, contig, del_start, del_end, buffer_size):
                continue
            
            # Reserve region
            reserve_region(reserved_regions, contig, del_start, del_end, buffer_size)
            
            # Store the plan
            sv_plans.append({
                'type': 'Deletion',
                'contig': contig,
                'start': del_start,  # 0-based
                'end': del_end,      # 0-based inclusive
                'size': size
            })
            
            placed += 1
        
        if placed < num_copies:
            print(f"Warning: Only placed {placed}/{num_copies} deletions of size {size}")
    
    # Plan insertions
    print("Planning insertions...")
    for size in sizes:
        placed = 0
        attempts = 0
        max_attempts = 10000
        
        while placed < num_copies and attempts < max_attempts:
            attempts += 1
            
            # Pick contig (weighted by length)
            contig = random.choices(contig_names,
                                   weights=[contig_lengths[h] for h in contig_names])[0]
            contig_len = contig_lengths[contig]
            
            # Pick insertion position
            ins_pos = random.randint(0, contig_len)
            
            # Check if position is available
            if check_buffer_overlap(reserved_regions, contig, ins_pos, ins_pos, buffer_size):
                continue
            
            # Reserve position
            reserve_region(reserved_regions, contig, ins_pos, ins_pos, buffer_size)
            
            # Generate random sequence for insertion
            random_seq = generate_random_sequence(size)
            
            # Store the plan
            sv_plans.append({
                'type': 'Insertion',
                'contig': contig,
                'position': ins_pos,  # 0-based
                'size': size,
                'sequence': random_seq
            })
            
            placed += 1
        
        if placed < num_copies:
            print(f"Warning: Only placed {placed}/{num_copies} insertions of size {size}")
    
    print(f"Successfully planned {len(sv_plans)} SVs")
    return sv_plans

def apply_svs_and_create_truth(original_contigs, sv_plans):
    """
    Apply structural variants and create truth table.
    
    Key insight: We must apply mutations in a specific order to maintain
    coordinate accuracy. Process from RIGHT to LEFT so earlier positions
    aren't affected by later modifications.
    
    Truth table coordinates are ALL relative to the ORIGINAL reference genome.
    """
    # Work on a copy
    mutated_contigs = {h: seq for h, seq in original_contigs.items()}
    truth_records = []
    
    # Group plans by contig
    operations_by_contig = defaultdict(list)
    
    # Convert plans to operations
    for plan in sv_plans:
        if plan['type'] == 'Deletion':
            operations_by_contig[plan['contig']].append({
                'op_type': 'deletion',
                'position': plan['start'],
                'end_position': plan['end'],
                'size': plan['size'],
                'original_plan': plan
            })
        elif plan['type'] == 'Insertion':
            operations_by_contig[plan['contig']].append({
                'op_type': 'insertion',
                'position': plan['position'],
                'size': plan['size'],
                'sequence': plan['sequence'],
                'original_plan': plan
            })
    
    # Process each contig independently
    for contig_name in sorted(operations_by_contig.keys()):
        ops = operations_by_contig[contig_name]
        
        # Sort operations: deletions before insertions at same position, then by position descending
        # Process from RIGHT to LEFT so earlier positions aren't affected
        ops.sort(key=lambda x: (x['position'], 0 if x['op_type'] == 'deletion' else 1), reverse=True)
        
        print(f"\nProcessing {len(ops)} operations on contig '{contig_name}'")
        
        for op in ops:
            if op['op_type'] == 'deletion':
                # Deletion: remove sequence
                pos = op['position']
                end_pos = op['end_position']
                
                print(f"  Deleting {end_pos - pos + 1}bp at position {pos} (1-based: {pos + 1})")
                
                # Apply deletion
                mutated_contigs[contig_name] = (
                    mutated_contigs[contig_name][:pos] + 
                    mutated_contigs[contig_name][end_pos + 1:]
                )
                
                # Record truth entry (ORIGINAL reference coordinates)
                truth_records.append({
                    'contig': contig_name,
                    'start_1based': pos + 1,
                    'end_1based': end_pos + 1,
                    'type': 'Deletion',
                    'size': op['size'],
                    'description': f"Deletion of {op['size']}bp"
                })
                
            elif op['op_type'] == 'insertion':
                # Insertion: add random sequence
                pos = op['position']
                seq = op['sequence']
                
                print(f"  Inserting {len(seq)}bp at position {pos} (1-based: {pos + 1})")
                
                # Apply insertion
                mutated_contigs[contig_name] = (
                    mutated_contigs[contig_name][:pos] + 
                    seq + 
                    mutated_contigs[contig_name][pos:]
                )
                
                # Record truth entry (ORIGINAL reference coordinates)
                truth_records.append({
                    'contig': contig_name,
                    'start_1based': pos + 1,
                    'end_1based': pos + 1,  # Insertion point
                    'type': 'Insertion',
                    'size': op['size'],
                    'description': f"Insertion of {op['size']}bp random sequence"
                })
    
    return mutated_contigs, truth_records

# ------------------ ADDED: VCF writer ------------------

def write_vcf(path, original_contigs, sv_plans):
    """
    Write a simple SV VCF (caller-like) from the *planned* SVs (original coordinates).

    Changes from previous version:
    - REF now contains the actual reference base (A/C/G/T)
    - ALT:
        * <DEL> for deletions (unchanged)
        * inserted sequence for insertions
    """
    contig_order = list(original_contigs.keys())
    contig_len = {c: len(original_contigs[c]) for c in contig_order}
    dt = datetime.now().strftime("%Y%m%d")

    with open(path, "w") as f:
        f.write("##fileformat=VCFv4.2\n")
        f.write(f"##fileDate={dt}\n")
        f.write("##source=SVMutator_InDel\n")
        f.write('##ALT=<ID=DEL,Description="Deletion">\n')
        f.write('##INFO=<ID=SVTYPE,Number=1,Type=String,Description="Type of structural variant">\n')
        f.write('##INFO=<ID=END,Number=1,Type=Integer,Description="End position of the variant">\n')
        f.write('##INFO=<ID=SVLEN,Number=1,Type=Integer,Description="Length of SV (negative for DEL)">\n')
        f.write('##INFO=<ID=IMPRECISE,Number=0,Type=Flag,Description="Imprecise structural variation">\n')
        for c in contig_order:
            f.write(f"##contig=<ID={c},length={contig_len[c]}>\n")
        f.write("#CHROM\tPOS\tID\tREF\tALT\tQUAL\tFILTER\tINFO\n")

        for idx, plan in enumerate(sv_plans, start=1):
            sv_id = f"SV{idx:05d}"
            chrom = plan["contig"]
            qual = "."
            flt = "PASS"

            if plan["type"] == "Deletion":
                pos = plan["start"] + 1
                end = plan["end"] + 1
                svlen = -plan["size"]

                # REF = reference base at deletion start
                ref = original_contigs[chrom][plan["start"]]

                alt = "<DEL>"
                info = f"SVTYPE=DEL;END={end};SVLEN={svlen}"
                f.write(f"{chrom}\t{pos}\t{sv_id}\t{ref}\t{alt}\t{qual}\t{flt}\t{info}\n")

            elif plan["type"] == "Insertion":
                pos = plan["position"] + 1
                end = pos
                svlen = plan["size"]

                # REF = reference base at insertion position
                ref = original_contigs[chrom][plan["position"]]

                # ALT = inserted sequence
                alt = plan["sequence"]

                info = f"SVTYPE=INS;END={end};SVLEN={svlen}"
                f.write(f"{chrom}\t{pos}\t{sv_id}\t{ref}\t{alt}\t{qual}\t{flt}\t{info}\n")

            else:
                raise ValueError(f"Unknown SV type in plan: {plan['type']}")

    print(f"Wrote VCF: {path} ({len(sv_plans)} records)")

    """
    Write a simple SV VCF (caller-like) from the *planned* SVs (original coordinates).

    - Deletion:  ALT=<DEL>, POS=start, END=end, SVLEN=-size
    - Insertion: ALT=<INS>, POS=position, END=position, SVLEN=+size

    REF is written as 'N' (common for symbolic SVs when base context is not provided).
    """
    contig_order = list(original_contigs.keys())
    contig_len = {c: len(original_contigs[c]) for c in contig_order}
    dt = datetime.now().strftime("%Y%m%d")

    with open(path, "w") as f:
        f.write("##fileformat=VCFv4.2\n")
        f.write(f"##fileDate={dt}\n")
        f.write("##source=SVMutator_InDel\n")
        f.write('##ALT=<ID=DEL,Description="Deletion">\n')
        f.write('##ALT=<ID=INS,Description="Insertion">\n')
        f.write('##INFO=<ID=SVTYPE,Number=1,Type=String,Description="Type of structural variant">\n')
        f.write('##INFO=<ID=END,Number=1,Type=Integer,Description="End position of the variant">\n')
        f.write('##INFO=<ID=SVLEN,Number=1,Type=Integer,Description="Length of SV (negative for DEL)">\n')
        f.write('##INFO=<ID=IMPRECISE,Number=0,Type=Flag,Description="Imprecise structural variation">\n')
        for c in contig_order:
            f.write(f"##contig=<ID={c},length={contig_len[c]}>\n")
        f.write("#CHROM\tPOS\tID\tREF\tALT\tQUAL\tFILTER\tINFO\n")

        # Deterministic IDs
        for idx, plan in enumerate(sv_plans, start=1):
            sv_id = f"SV{idx:05d}"
            chrom = plan["contig"]
            ref = "N"
            qual = "."
            flt = "PASS"

            if plan["type"] == "Deletion":
                pos = plan["start"] + 1
                end = plan["end"] + 1
                svlen = -plan["size"]
                alt = "<DEL>"
                info = f"SVTYPE=DEL;END={end};SVLEN={svlen}"
                f.write(f"{chrom}\t{pos}\t{sv_id}\t{ref}\t{alt}\t{qual}\t{flt}\t{info}\n")

            elif plan["type"] == "Insertion":
                pos = plan["position"] + 1
                end = pos
                svlen = plan["size"]
                alt = "<INS>"
                info = f"SVTYPE=INS;END={end};SVLEN={svlen}"
                f.write(f"{chrom}\t{pos}\t{sv_id}\t{ref}\t{alt}\t{qual}\t{flt}\t{info}\n")

            else:
                raise ValueError(f"Unknown SV type in plan: {plan['type']}")

    print(f"Wrote VCF: {path} ({len(sv_plans)} records)")

# -------------------------------------------------------

def main(args):
    """Main execution"""
    if not Path(args.input).exists():
        raise FileNotFoundError(f"Input file not found: {args.input}")
    
    print("="*60)
    print("Structural Variant Mutator - Insertions & Deletions")
    print("="*60)
    
    # Load original reference genome
    original_contigs = read_fasta(args.input)
    
    # Plan all SVs (no mutations applied yet)
    sv_plans = plan_all_svs(
        original_contigs=original_contigs,
        sizes=SV_SIZES,
        num_copies=NUM_COPIES,
        buffer_size=args.buffer_size,
        seed=args.seed
    )
    
    # Apply mutations and create truth table
    mutated_contigs, truth_records = apply_svs_and_create_truth(
        original_contigs=original_contigs,
        sv_plans=sv_plans
    )
    
    # Write outputs
    write_fasta(args.output, mutated_contigs)
    write_tsv(args.tsv, truth_records)

    # ADDED: write VCF output (planned SVs, original coordinates)
    write_vcf(args.vcf, original_contigs, sv_plans)
    
    # Summary
    print("\n" + "="*60)
    print("Summary:")
    print(f"  Total SVs applied: {len(sv_plans)}")
    print(f"  Deletions: {sum(1 for p in sv_plans if p['type'] == 'Deletion')}")
    print(f"  Insertions: {sum(1 for p in sv_plans if p['type'] == 'Insertion')}")
    print(f"  Truth records: {len(truth_records)}")
    print("="*60)
    print("\nNote: All coordinates in the truth table are relative to the")
    print("      ORIGINAL reference genome for validation purposes.")
    print("="*60)

if __name__ == "__main__":
    p = argparse.ArgumentParser(
        description="Generate structural variants (insertions & deletions) in a genome"
    )
    p.add_argument("--input", default=INPUT_FASTA, 
                   help="Input FASTA file")
    p.add_argument("--output", default=OUTPUT_FASTA,
                   help="Output mutated FASTA file")
    p.add_argument("--tsv", default=MUTATION_TSV,
                   help="Output truth table TSV file")
    p.add_argument("--vcf", default=OUTPUT_VCF,
                   help="Output VCF file with SV records")
    p.add_argument("--buffer-size", type=int, default=BUFFER_SIZE,
                   help="Buffer size to avoid overlapping SVs")
    p.add_argument("--seed", type=int, default=SEED,
                   help="Random seed for reproducibility")
    
    args = p.parse_args()
    main(args)
```
