## Supplementary material for "Easy-to-use whole-genome sequencing workflows and standardized practices to uncover hidden genetic variation in *Synechocystis* PCC 6803 wild-type and knock-out strains": S9 File

### Plasmids, Primer and Strains used

#### Plasmids

| Name | Origin | Used for |
|---|---|---|
| **Δgnd** | This study | Generation of Δgnd_3 mutant |
| **pSSR:gGND** | This study | Plasmid-based complementation under an inducible rhamnose promoter of Δgnd strains |
| **pSS_Pgnd:GND** | This study | Plasmid-based complementation under the native promoter of Δgnd strains |
| **pBS_GND_comp** | This study | Genome-based complementation of Δgnd strains |
| **pSSR** | This study | pSHDY (Behle et al., 2020) based plasmid containing a rhamnose-inducible promoter, stable in Synechocystis PCC 6803 |
| **pSSR:C_N-6His-Tag** | This study | pSSR-based plasmid containing restriction sites to add genes under the rhamnose inducible promoter and fusing them to either an N or C-terminal 6-His tag |

#### Primer

| Name | Sequence | Used for |
|---|---|---|
| **MT_P0046_pSSRV_Seq_F** | GTTGCCAATGGCCCATTTTCC | Check if strains contain pSSR plasmids, Sequencing pSSR-based plasmids |
| **SDHY_rev** | AGCTCCATAGGCCGCTTT | Check if strains contain pSSR plasmids, Sequencing pSSR-based plasmids |
| **pSSR-GND-Synechocystis-6His.FOR** | GAGTAGTGGAGGTTACCATGTGCAATTTAACGTAGCTATTATGACTAAGCG | Cloning the gnd gene into pSSR:C_N-6His-Tag via Gibson assembly |
| **pSSR-GND-Synechocystis-6His.REV** | CCTGAAAATACAGGTTTTCGGATCCATCAAGCCATTCCGTGTGGAAA | Cloning the gnd gene into pSSR:C_N-6His-Tag via Gibson assembly |
| **Gnd_For** | tatgctcttctgctcctgcatcaatcaagccattccgtg | Cloning 130bp upstream + the gnd gene into pSSR-N |
| **Gnd_rev** | cccagcctgcgcgatgtctagatcgccgatgaacctagc | Cloning 130bp upstream + the gnd gene into pSSR-N |
| **MB08_B9** | ATGGCCCATTTTCCTGTCAGTAACGAG | Check if strains contain pSSR plasmids, Sequencing pSSR-based plasmids |
| **MB08_C1** | TTTGCTTCCAGATGTATGCTCTTCTGC | Check if strains contain pSSR plasmids, Sequencing pSSR-based plasmids |

| | | |
|---|---|---|
| **MT_P0038_2xHis_A_F** | TATGCATCACCATCACCATCACGGCGCTAGCCCCGGGGAAAACCTGTATTTTCAGGGCG | First fragment to create pSSR:C_N-6His-Tag via ligation into pSSR-N |
| **MT_P0039_2xHis_A_R** | GATCCGCCCTGAAAATACAGGTTTTCCCCGGGGCTAGCGCCGTGATGGTGATGGTGATGCA | First fragment to create pSSR:C_N-6His-Tag via ligation into pSSR-N |
| **MT_P0040_2xHis_B_F** | GATCCGAAAACCTGTATTTTCAGGGCGGCGCTAGCCCCGGGCATCACCATCACCATCACTAAAGG | Second fragment to create pSSR:C_N-6His-Tag via ligation into pSSR-N |
| **MT_P0041_2xHis_B_R** | CCTTTAGTGATGGTGATGGTGATGCCCGGGGCTAGCGCCGCCCTGAAAATACAGGTTTTCG | Second fragment to create pSSR:C_N-6His-Tag via ligation into pSSR-N |
| **MT_P0029_dGND_gibson_R** | AGGGAACAAAAGCTGGAGCTAGCTTGCCACCCTCTGACTG | Used to amplify the genomic region ~230bp up and downstream of the gnd locus from the Δgnd_zwf399+A mutant and cloning it into the pBS plasmid |
| **MT_P0030_dGND_gibson_F** | CTATAGGGCGAATTGGGTACAAGGAATTGCTTTGACCCACGAAG | Used to amplify the genomic region ~230bp up and downstream of the gnd locus from the Δgnd_zwf399+A mutant and cloning it into the pBS plasmid |
| **MT_P0028_dGND_rescue_R** | AAGGAATTGCTTTGACCCACGAAG | Amplify the genomic region of *gnd* to check segregation |
| **MT_P0027_dGND_rescue_F** | AGCTTGCCACCCTCTGACTG | Amplify the genomic region of *gnd* to check segregation |
| **MT_204_pSS_Pgnd-GND_Seq.FOR** | tgatgttacccgagagcttgg | Amplify the insert in the pSDHY based complementation plasmid *Pgnd:gnd* |
| **MT_205_pSS_Pgnd-GND_Seq.REV** | gcggcgggcaagtacg | Amplify the insert in the pSDHY based complementation plasmid *Pgnd:gnd* |

### Strains

| Short name | Full Name | Origin | Description |
|---|---|---|---|
| **Δgnd_zwf399+A** | Δgnd::EmR zwf+399insA | (Chen et al., 2016a) | Replacement of the gnd gene with an Erythromycin |

| | | | |
|---|---|---|---|
| | | | resistance cassette, additional frameshift mutation in ZWF, |
| **Δgnd::gnd_zwf399+A** | Δgnd::gnd,SpecR zwf+399insA | This Study | Complementation of the Δgnd_zwf399+A strain by reintroducing the gnd gene, followed by a Spectinomycin resistance cassette into the original locus |
| **Δgnd_zwf399+A_ pSSR:GND** | Δgnd::EmR pSSR:gnd zwf+399insA | This Study | Complementation of the Δgnd_zwf399+A strain by introducing a stable plasmid containing the gnd gene under an rhamnose-inducible promoter |
| **Δgnd_zwf399+A_pSS_ Pgnd:GND** | Δgnd::EmR pSS_Pgnd:GND zwf+399insA | This Study | Complementation of the Δgnd_zwf399+A strain by introducing a stable plasmid containing the gnd gene and 130bp upstream of the gene containing the native gnd promoter |
| **Δgnd_3** | Δgnd::EmR | This Study | Replacement of the gnd gene with an Erythromycin resistance cassette |
| **Δgnd_3::gnd** | Δgnd::gnd,SpecR | This Study | Complementation of the Δgnd_3 strain by reintroducing the gnd gene, followed by a Spectinomycin resistance cassette into the original locus |
| **Δgnd_3_pSS_ Pgnd:GND** | Δgnd::EmR pSS_Pgnd:GND | This Study | Complementation of the Δgnd_3 strain by introducing a stable plasmid containing the gnd gene and 130bp upstream of the gene containing the native gnd promoter |
| **Δzwf** | Δzwf::CmR | (Chen et al., 2016a) | Replacement of the gnd gene with a Chloramphenicol resistance cassette, additional frameshift mutation in ZWF, |

| | | |
|---|---|---|
| **WT_Kassel** | Originated from the McIntosh Lab | Synechocystis sp. PCC 6803 GT-Williams/ GT-T strain (according to (Koskinen et al., 2023), list of strain-specific mutations in the supplementary information, all knockout mutants used are based on this strain |
| **WT-Freiburg** | | Kindly gifted by Annegret Wilde, Freiburg, Germany, a list of strain-specific mutations in the supplementary information |
| **WT_Uppsala** | | Glucose tolerant non motile strain kindly gifted by Pia Lindberg, Uppsala, Sweden, a list of strain-specific mutations in the supplementary information |
| **WT_Stockholm** | | Glucose tolerant non motile strain kindly gifted by Paul Hudson Lab, Stockholm, Sweden, list of strain-specific mutations in the supplementary information |

### Strain validation

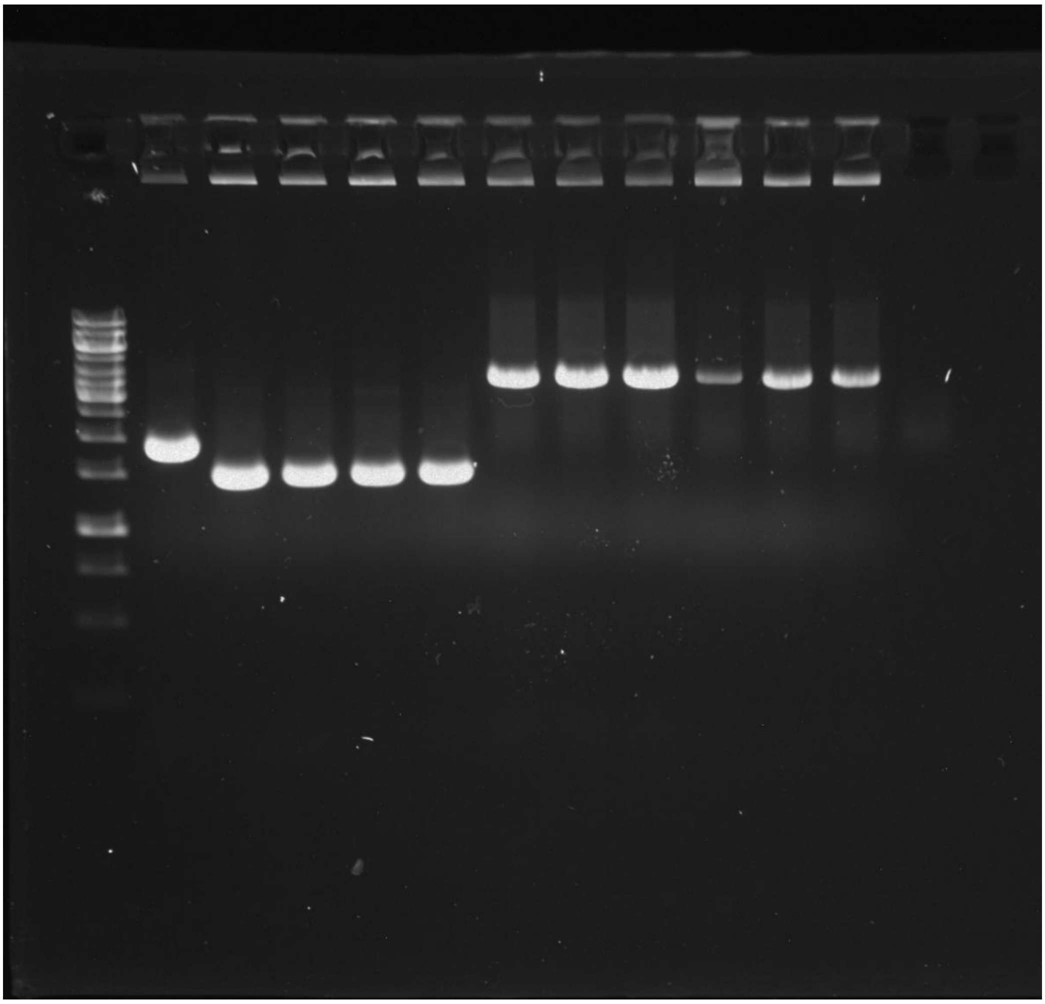

**Agarose Gel Electrophoresis of PCR samples to control segregation of knock-out and complemented strains**.

PCR with primer MTP0027 and MTP0028 that amplify the genomic region of *gnd*, expected bands are WT: 1926bp, Δ*gnd* : 1486bp, Δ*gnd*::gnd: 3482bp.

Lane 1: 1kb GeneRuler (Marker), Lane 2: WT, Lane 3:*Δgnd_zwf*$^{399+A}$, Lane 4-6: *Δgnd_3*, Lane 7-9: *Δgnd::gnd_zwf*$^{399+A}$, Lane 10-12: *Δgnd_3::gnd*

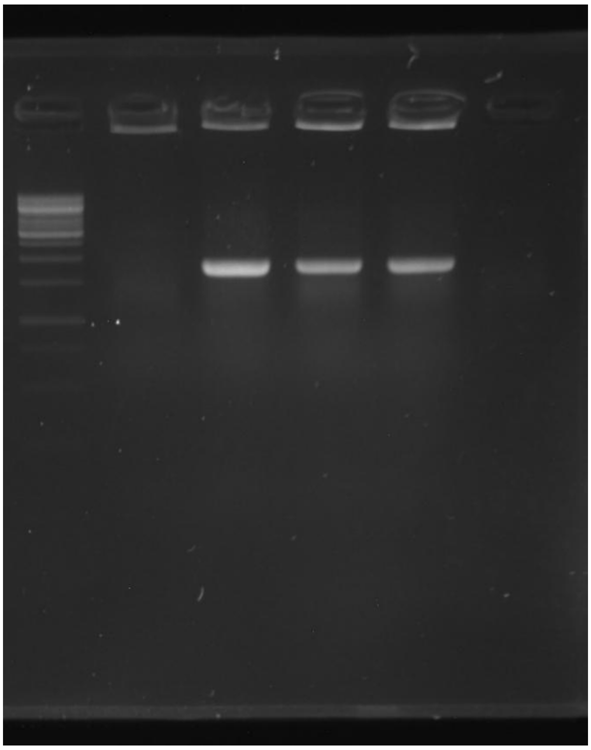

**Agarose Gel Electrophoresis of PCR samples to control the presence of the complementation plasmid *Pgnd:gnd*.**

PCR with primer MT_204_pSS_Pgnd-GND_Seq.FOR and MT_205_pSS_Pgnd-GND_Seq.REV that amplify the insert in *Pgnd:gnd* WT: no band, *Pgnd:gnd* 1834bp.
Lane 1: 1kb GeneRuler (Marker), Lane 2: WT, Lane 3-5:*Δgnd_3 Pgnd:gnd*
