## Supplementary material for "Easy-to-use whole-genome sequencing workflows and standardized practices to uncover hidden genetic variation in *Synechocystis* PCC 6803 wild-type and knock-out strains": S10 Figure

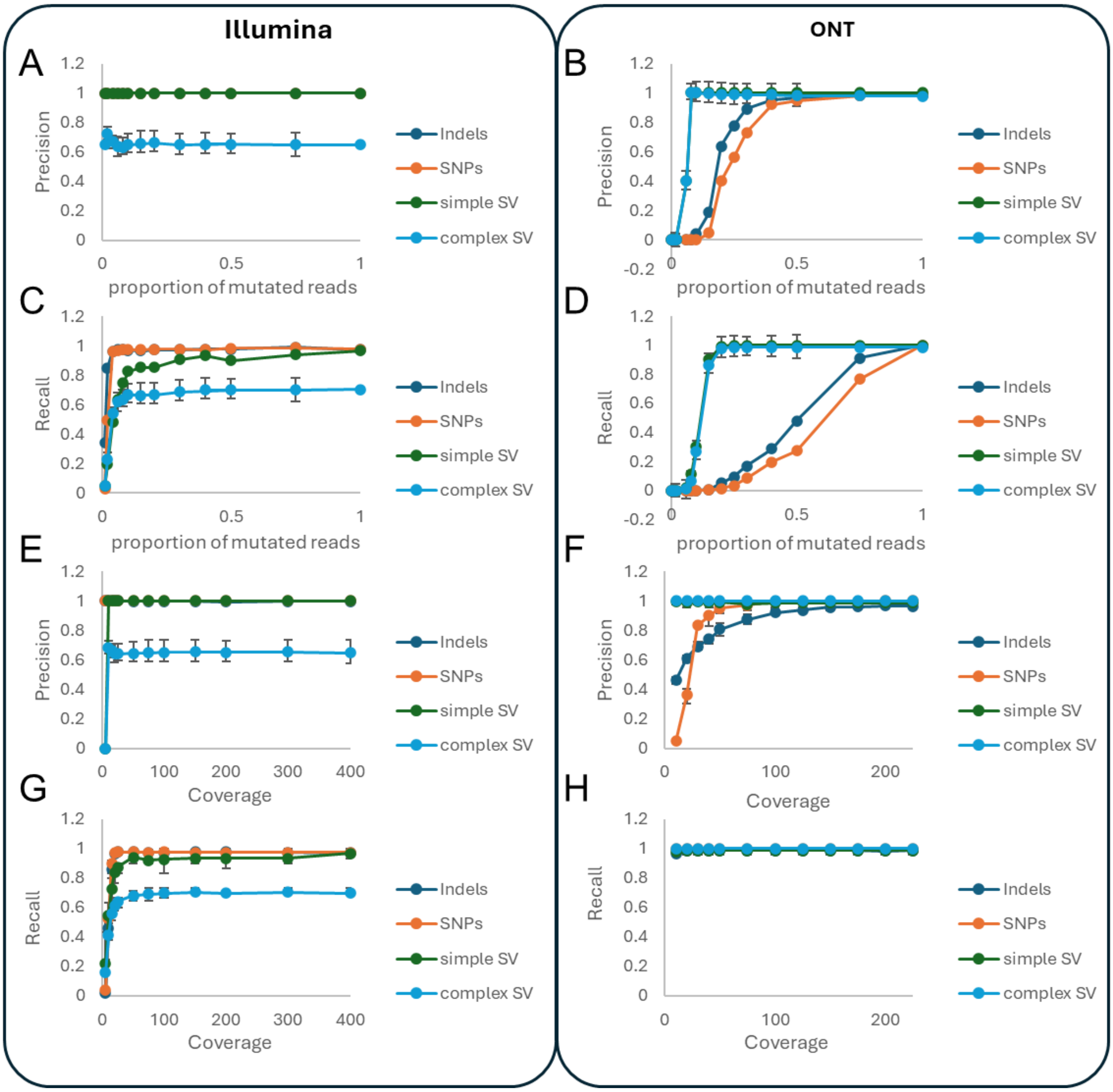


**Fig. S 10 Variant call pipelines were tested using simulated reads from intentionally altered genomes (for details, see the methods section).**

Variant call pipelines were tested using simulated reads from intentionally altered genomes (see the methods section for details). A, C) Precision and recall of variants called from simulated reads with changing ratios of alternative reads representing non-segregated mutants for Illumina sequencing (coverage 500x). B, D) Precision and recall of variants called from simulated reads with changing ratios of alternative reads representing non-segregated mutants for ONT sequencing (coverage 225x). E, G) precision and recall of variants called from simulated Illumina reads with different coverages (average coverage of 5, 10, 15, 20, 25, 50, 75, 100, 150,200, 300,400). F, H) precision and recall of variants called from simulated ONT (Oxford Nanopore Technology) reads with different coverages (average coverage of 10, 20, 30, 40, 50, 75, 100, 125, 150, 175, 200, 225).
