## Supplementary material for "Easy-to-use whole-genome sequencing workflows and standardized practices to uncover hidden genetic variation in *Synechocystis* PCC 6803 wild-type and knock-out strains": S11 Figure

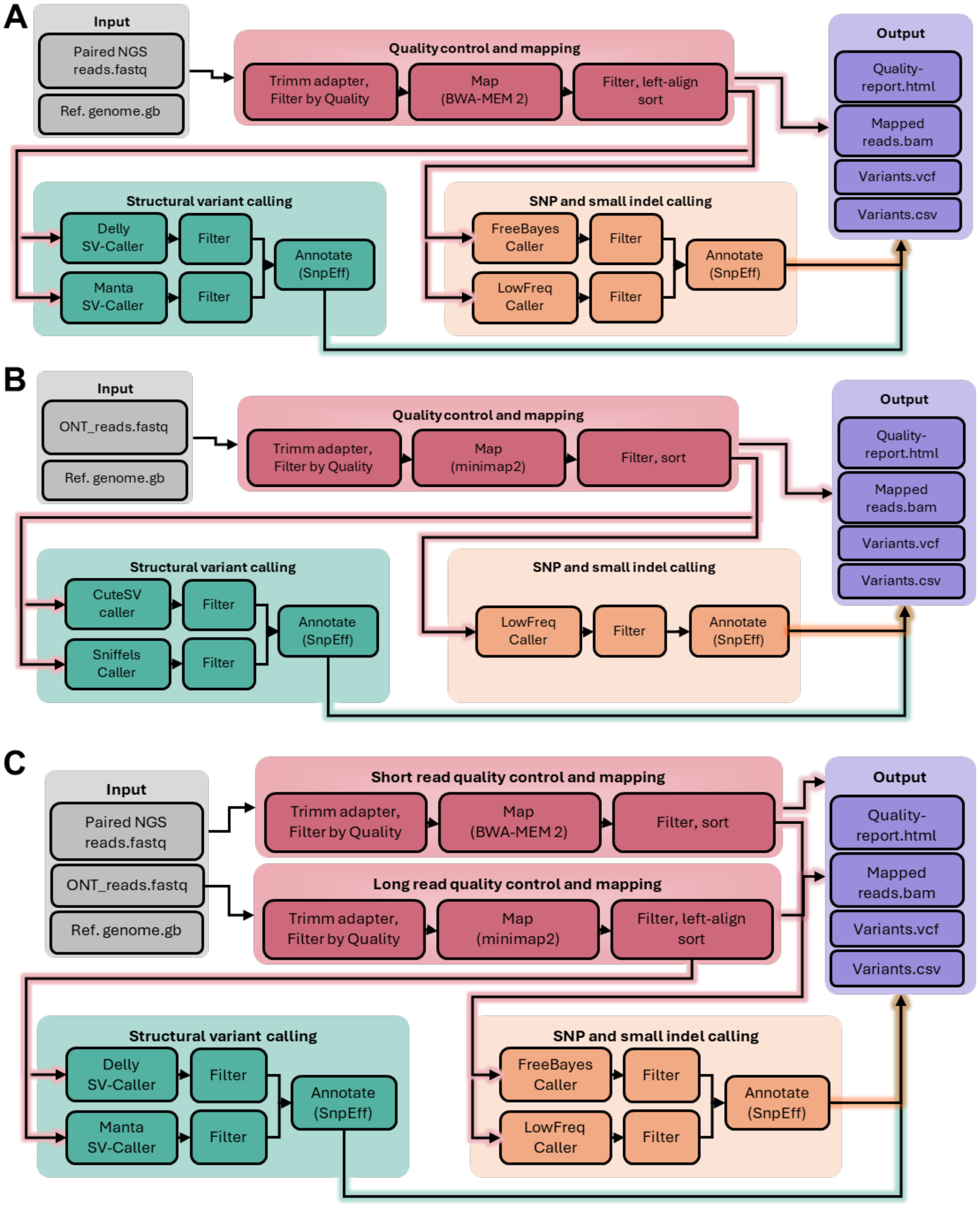


**Fig. S 11: Schematic but more detailed overview of the created galaxy workflows**

A) Workflow for short-read paired Illumina sequencing data. B) Workflow for long-read ONT (Oxford Nanopore technology) sequencing reads. C) Hybrid workflow were paired end Illumina reads are used for SNP and small indel calling, while long reads are used for structural variant detection.
